# Global cellular response to chemical perturbation of PLK4 activity and abnormal centrosome number

**DOI:** 10.1101/2021.06.25.449796

**Authors:** Johnny M Tkach, Reuben Philip, Amit Sharma, Jonathan Strecker, Daniel Durocher, Laurence Pelletier

## Abstract

Centrosomes act as the main microtubule organizing centre (MTOC) in metazoans. Centrosome number is tightly regulated by limiting centriole duplication to a single round per cell cycle. This control is achieved by multiple mechanisms, including the regulation of the protein kinase PLK4, the most upstream facilitator of centriole duplication. Altered centrosome numbers in mouse and human cells cause p53-dependent growth arrest through poorly defined mechanisms. Recent work has shown that the E3 ligase TRIM37 is required for cell cycle arrest in acentrosomal cells. To gain additional insights into this process, we undertook a series of genome-wide CRISPR/Cas9 screens to identify factors important for growth arrest triggered by treatment with centrinone B, a selective PLK4 inhibitor. We found that TRIM37 is a key mediator of growth arrest after partial or full PLK4 inhibition. Interestingly, PLK4 cellular mobility decreased in a dose-dependent manner after centrinone B treatment. In contrast to recent work, we found that growth arrest after PLK4 inhibition correlated better with PLK4 activity than with mitotic length or centrosome number. These data provide insights into the global response to changes in centrosome number and PLK4 activity and extend the role for TRIM37 in regulating the abundance, localization and function of centrosome proteins.

## INTRODUCTION

The centrosome is a multi-protein complex that is the major MTOC of metazoan cells influencing microtubule-based processes such as cell division, ciliogenesis, signaling and cell motility [1]. Each centrosome consists of two microtubule-based centrioles surrounded by pericentriolar material (PCM) [1]. Each cell inherits a single centrosome from the previous cell cycle and subsequent centriole duplication is restricted to a single round of replication [2]. Centriole duplication initiates at the G1/S transition and is largely a stepwise pathway dependent on the upstream kinase PLK4 [2]. PLK4 is a low-abundance protein [3] that is recruited around the mother centriole [4, 5]. PLK4 kinase activity is regulated by interaction with STIL and CEP85 resulting in the recruitment of SASS6 to form the cartwheel of the nascent daughter centriole followed by the recruitment of centriole elongation factors [6–13]. PLK4 functions as a homodimer and autophosphorylates itself in trans to generate a phosphodegron sequence that limits its abundance. This sequence is recognized by SCF^β-TrCP^ and targets PLK4 for ubiquitin-mediated proteolysis [14–16].

PLK4 misregulation is often associated with pathological states. Centrosome amplification is a hallmark of tumour cells [17] and may play a role in generating chromosome instability [18, 19] and promoting cell invasiveness [20]. In cell culture and mouse models, overexpression, inhibition or deletion of PLK4 results in p53-dependent arrest [21–29]. A series of CRISPR/Cas9 screens identified the p53 pathway members, p53, p21(CDKN1A), 53BP1 and USP28, [22, 24, 30], and the E3-ligase TRIM37 as component of this response [24, 30]. A recent screen for mediators of supernumerary centrosome-induced arrest identified PIDDosome/p53 and placed the distal appendage protein ANKRD26 within this pathway [31, 32].

TRIM37 is an E3 ligase that has been associated with a myriad of cellular functions including gene expression [33], peroxisome maturation [34], various signaling pathways [35–38] and centriole biology [39–42]. There is no consensus on how TRIM37 mediates these functions since its activities have been linked to mono-ubiquitination [33, 34], poly-ubiquitination [35, 37, 42, 43] and E3-independent functions [38] that result in changes in protein activity [33, 38], localization [30, 40, 41] and abundance [34, 35, 37, 42, 43]. A number of centrosome-related TRIM37 functions have been described. In the absence of TRIM37 a collection of centriole proteins such as CNTROB, PLK4, CETN1/2, CP110 accumulate to form aberrant assemblies referred to as condensates or ‘Cenpas’ (Centriolar protein assemblies) [40, 41]. These structures are dependent on the presence of CNTROB, but it is unclear why TRIM37 might suppress their formation in normal cells. TRIM37 is part of the 17q23 amplicon present in approximately 18% of breast cancer tumours [44] and overexpression of TRIM37 in these lines renders them sensitive to the PLK4 inhibitor centrinone [42, 43]. TRIM37 interacts with PLK4 and CEP192 [43]. Although TRIM37 can promote ubiquitination of PLK4, it is distinct from SCF^β−TRCP^ modification since it does not result in changes to PLK4 abundance [43]. In contrast, transiently overexpressed TRIM37 leads to CEP192 ubiquitination and its subsequent degradation [42, 43]. In this model of TRIM37 function, PLK4-nucleated condensates consisting of PCM components facilitate mitosis in the absence of TRIM37 and the overexpression of TRIM37 decreases the cellular levels of CEP192 rendering cells sensitive to the loss of centrioles.

Here, we sought to determine how growth arrest is initiated in response to alterations in PLK4 activity and centrosome number. Using the specific PLK4 inhibitor, centrinone B we modulated PLK4 activity to generate supernumerary centrosomes or centrosome loss and performed genome-wide chemical genetic screens in RPE-1 and A375 cells. Our screens identified distinct pathways mediating the response to partial and full PLK4 inhibition. Intriguingly, TRIM37 was required for growth when PLK4 was partially or fully inhibited but was dispensable for arrest triggered by PLK4 overexpression. Moreover, TRIM37 growth arrest activity was partially independent of its E3 ligase activity. These results highlight the complex role of TRIM37 and regulators of its function in the control of centrosome number homeostasis.

## RESULTS

### Centrinone B induces concentration-dependent changes in centrosome number

Centrinone is a PLK4-specific inhibitor that is often used to study the cellular response to centrosome loss [23]. Another PLK4 inhibitor, CFI-400945, induces both centrosome amplification, or loss, in a concentration-dependent manner [45]. Since CFI-400945 also inhibits other mitotic kinases such as AURKB [46] it is unclear if its reported phenotypes are due to effects on PLK4, mitosis, or both. Centrinone and centrinone B are both potent inhibitors of PLK4 that show selectivity over the Aurora-family kinases [23]. We used centrinone B for our experiments since it shows an even greater selectivity over the Aurora kinases compared to centrinone [23, 46]. We treated RPE-1 cells with 200 and 500 nM centrinone B and found that cell growth was greatly inhibited at both centrinone B concentrations (Figure 1A). As expected, cell growth arrest was correlated with induction of p53 and p21 (Figure 1B and C and Figure 1 -- figure supplement 1A and B), resulting in the accumulation of cells with 1 N DNA content (Figure 1 – figure supplement 1C). We next imaged cells treated with centrinone B for 3 days to determine how centrosome number was affected as a function of centrinone B concentration. As expected, 500 nM centrinone B resulted in cells containing either a single or no centrosome, but cells treated with 200 nM failed to lose centrosomes and instead accumulated supernumerary centrosomes in approximately 50% of cells (Figure 1C and D, Figure 1- figure supplement 1B and D). Staining for a panel of centriole and centrosome markers revealed that these extra structures contained CEP135, CEP120, CETN2, glutamylated tubulin and could accumulate PCNT and CEP192 indicating that treatment with 200 nM centrinone B can induce amplification of *bona fide* centrosomes (Figure 1 – figure supplement 1E). Together, these data indicate that centrosomes can be amplified or lost using 200 or 500 nM centrinone B, respectively, and that both phenotypes result in p53-dependent G1 arrest.

**Figure 1.**
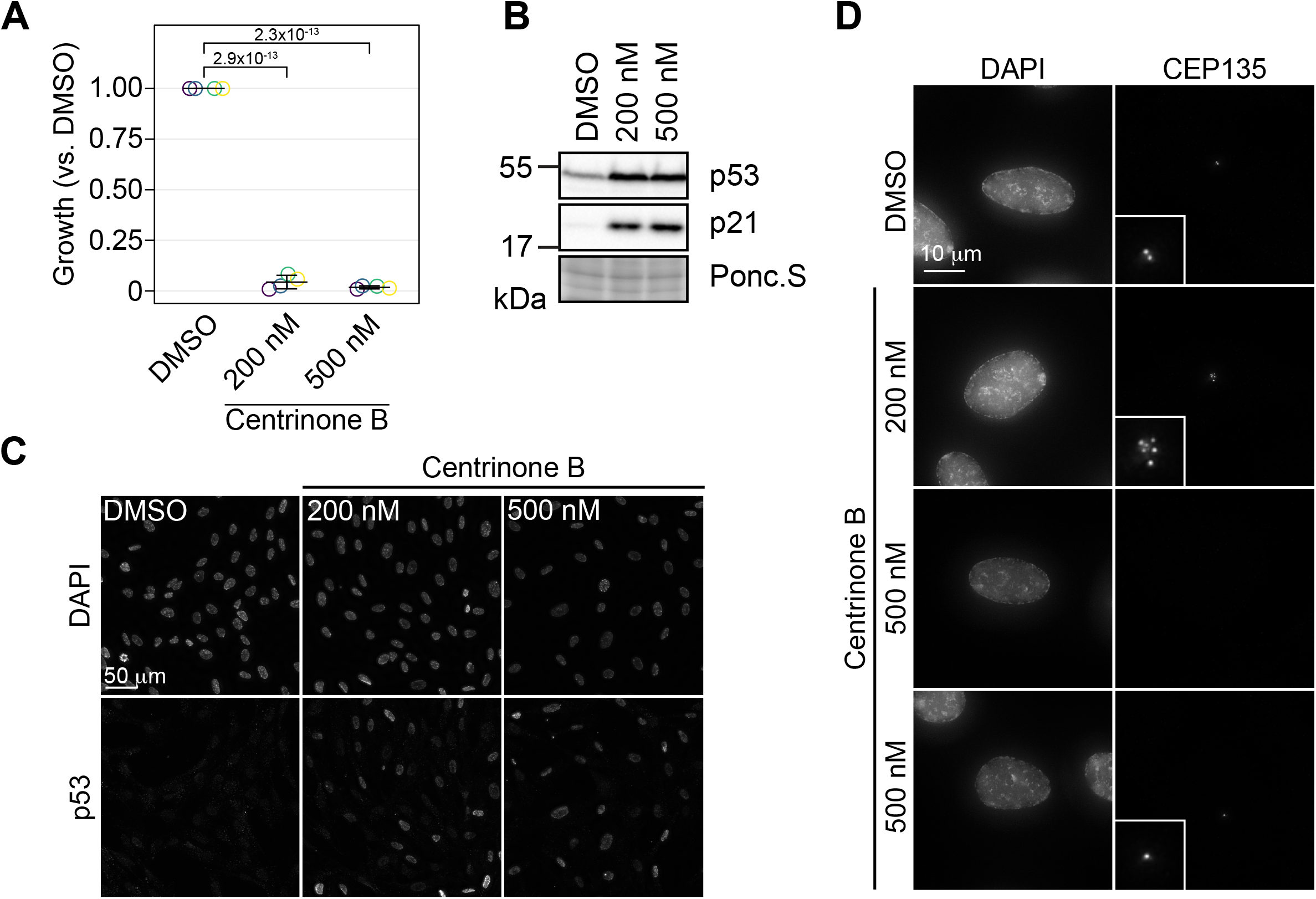
Concentration-dependent effect of centrinone B on centriole number. **(A)** RPE-1 cells were serially grown for 12 days and treated with DMSO or the indicated concentration of centrinone B. Relative cell number compared to a DMSO-treated control was determined and plotted. Three independent replicates plotted with mean with standard deviation shown. Significant *p*-values from Dunnett post-hoc test using ‘DMSO’ as control after one-way ANOVA shown. **(B)** RPE-1 cells were treated with DMSO or 200 or 500 nM centrinone B for 4 days and prepared for Western blot probing for the indicated proteins. Ponc.S indicates total protein. **(C)** RPE-1 cells were treated as in (B), fixed for immunofluorescence and stained for p53. **(D)** RPE-1 cells were treated as in (B), fixed for immunofluorescence and stained for CEP135. Examples of cells with no centrosomes or one centrosome are shown. Inset magnified 3x.

### Global cellular responses to abnormal centrosome number

To understand the mechanisms of centrinone-induced cell cycle arrest we performed genome-wide CRISPR screens in the presence of 200 nM and 500 nM centrinone B (Figure 2A) [47]. We reasoned that cell fitness at each centrinone B condition would require distinct sets of genes. Since loss of components in the p53 pathway itself would also increase fitness, we performed a parallel screen in the presence of Nutlin-3a, a small molecule that prevents the MDM2-mediated inactivation of p53 [48], allowing us to filter out core p53 pathway components. RPE-1 or A375 cells stably expressing Cas9 were infected with the TKOv1 lentiviral sgRNA library [47], selected, and subsequently treated with the indicated drug concentrations (Figure 2A). After growth for 21 d, cells were harvested and subjected to next-generation sequencing (NGS) and Model-based Analysis of Genome-wide CRISPR-Cas9 Knockout (MAGeCK) analysis [49]. Genes with a false discovery rate (FDR) of < 0.05 were considered hits for subsequent analyses. The combined screens to identify regulators of growth arrest using Nutlin-3a, 200 nM and 500 nM centrinone B yielded 91, 136 and 56 high-confidence hits that positively or negatively affected cell growth, respectively (Table S1).

**Figure 2.**
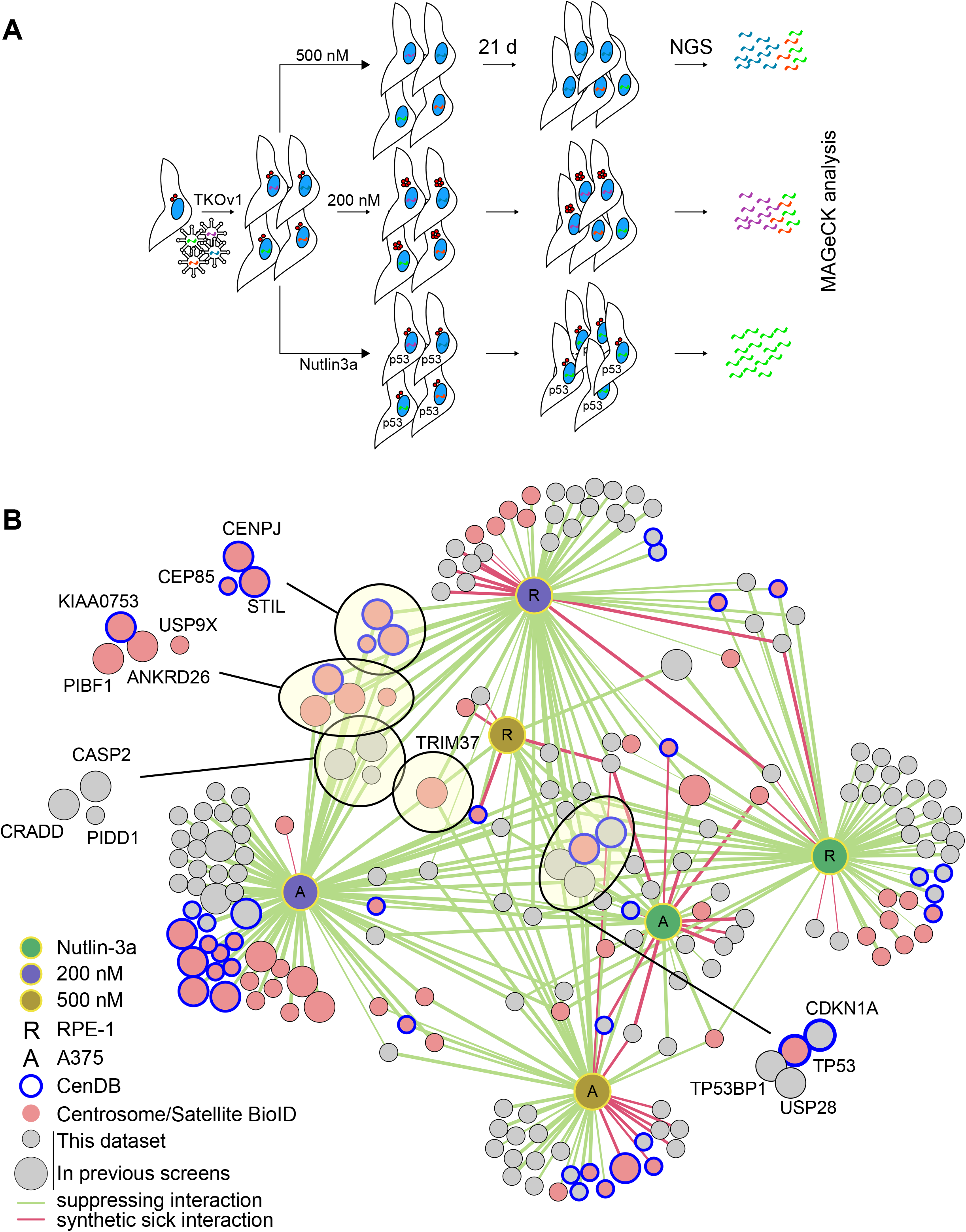
CRISPR/Cas9 screen to interrogate response to abnormal centrosome number. **(A)** Schematic outlining our screening procedures. Cells expressing Cas9 were infected with the TKOv1 genome-wide CRISPR sgRNA library and subsequently grown for 21 days in the presence of DMSO, 200 nM centrinone B, 500 nM centrinone B or 600 nM Nutlin-3a. Genomic DNA was prepared and sgRNA counts in each pool of cells was determined using NGS and analyzed using MAGeCK. Screens were performed in technical triplicate. **(B)** The significant hits (p < 0.05) from all screens were combined to form a network. Each unique cell and drug combination used for screening were set as hubs (i.e., RPE-1 200 nM centrinone B). All other nodes represent genes identified. Edges connect identified genes with a screening condition with edge weight inversely proportional to FDR. The general layout using the automated yFiles organic method from Cytoscape was preserved while individual nodes were manually arranged to facilitate visualization. Selected complexes and protein nodes are circled and highlighted. Except for the hubs, large nodes represent genes identified by previous PLK4 inhibition screens (see Table S1).

We created a network diagram to visualize the hits on a global scale (Figure 2B). Each unique cell line and condition act as the hubs (i.e., RPE-1, 200 nM centrinone B) while the hits from each condition are the remaining nodes. Each screen identified overlapping and distinct sets of genes, supporting our hypothesis that cells respond differently to each of the conditions tested (Figure 2B, Figure 2 – figure supplement 2A-D). The Nutlin-3a dataset contained core p53 pathway genes *TP53* and *CDKN1A* and genes coding for p53 regulators *TP53BP1* and *USP2*8, consistent with their role in promoting p53 transcriptional activity [50]. This dataset likely contains other mediators of the p53 pathway. The disruption of both *BAG6* and *EP300* increased fitness in Nutlin-3a. EP300 is an acetyltransferase that binds to and affects the acetylation of p53 while BAG6 modulates this acetylation event by EP300 [51, 52]. Likewise, inactivation of *AHR* and *ARNT*, which form a transcriptional complex activated by exogenous ligands, promoted growth when p53 is activated in RPE-1 cells, but not A375. ARNT was previously identified as a FRET interactor with p53 [53] and interacts with EP300 [54, 55]. Deletion of *TSC1/2*, that integrates p53 signalling with the mTOR pathway [56, 57] caused decreased fitness after p53 activation. We used Genemania [58] to further probe the pairwise physical interactions among the hits from the RPE-1 Nutlin-3a screen and generated a significantly enriched network (∼20-fold enrichment, *p* = 4.1×10^−31^). In this network, 27 of the 57 hits formed physical interactions with eight proteins forming complexes with p53 itself (Figure 2 – figure supplement 1E). Our high confidence hits from the Nutlin-3a screen identified known p53 pathway members and likely contains unknown regulators of this pathway that will warrant further characterization.

The 200 nM centrinone B screens (i.e. condition that produces supernumerary centrosomes) revealed a core set of 23 genes that suppressed the growth arrest in both cell lines. (Table S1, Figure 2 – figure supplement 2C). Notably, we identified the ANKRD26/CASP2/PIDD1/CRADD (PIDDosome) complex recently implicated in the response to supernumerary centrosomes [31, 32]. This set also included p53 pathway genes (*TP53*, *CDKN1A*, *TP53BP1* and *USP28*), centriole duplication factors (*CEP85*, *CENPJ*, *STIL*, *USP9X*), centriolar satellite proteins (*C2CD3*, *CEP350*, *KIAA0753*, *PIBF1*) and *TRIM37*. The A375 screen identified additional centriole components namely *CP110*, *CEP97*, *CEP135*, *SASS6*, *CEP76* and *PLK4*. Twenty-two of the hits from both cell lines combined also overlapped with the high-confidence hits from a similar screen that induced centrosome amplification by overexpressing PLK4 [31]. The overexpression screen also yielded the ANKRD26/PIDDOsome and some, but not all the centriole-associated genes, nor *TRIM37* (Figure 2 – figure supplement 2F).

We identified a total of 37 suppressors probing the response to centriole depletion (500 nM centrinone B), with five scoring in both cell lines (*TP53*, *CDKN1A*, *TP53BP1*, *USP28* and *TRIM37*). Our results are similar to previous screens aimed at identifying suppressors of growth arrest due to centrosome loss from PLK4 inhibition that identified 31, 41 and 27 genes respectively [22, 24, 30]. Four of the five common hits (*TP53*, *TP53BP1*, *USP28*, *CDKN1A*) correspond to core p53 pathway components and were the only genes identified by all screens performed to date (Figure 2 – figure supplement 2G). *CHD8* and *FBXO42* were previously identified in the response to centrosome loss [24] and also scored in our 200 nM and 500 nM centrinone B screens, respectively; however they also appeared in our Nutlin-3a hits suggesting that these genes might not be specific to centrosome biology. Indeed, both FBXO42 and CHD8 are known to negatively regulate p53 activity [59–61]. Two of the previous centriole loss screens also identified *TRIM37* [22, 30] which was unique among all the other hits since it was the only gene outside the p53 pathway that scored in both RPE-1 and A375 cells in both centrinone B concentrations. We therefore chose to study TRIM37 further.

### TRIM37 localizes near the centrosomes but is not required for centriole duplication

Our combined screens indicated that TRIM37 was required for growth arrest in response to PLK4 inhibition that results in either centrosome over-duplication or loss. Since TRIM37 has been implicated in centriole duplication [39], we determined if loss of TRIM37 affected centrosome number after treatment with centrinone B. RPE-1 cells were infected with viruses expressing two independent sgRNAs targeting *TRIM37* (Figure 3 – figure supplement 1A and B) and selected cell pools were treated with 200 and 500 nM centrinone B and assessed for centriole number (Figure 3A). As previously observed [39], we noted a small number of *TRIM37*-disrupted cells harboured extra centrioles in untreated cells. However, centriole amplification or loss after centrinone B treatment was not greatly affected (Figure 3A). To further characterize TRIM37, we created a *TRIM37*-disrutped RPE-1 clonal cell line using CRISPR/Cas9 (Figure 3 – figure supplement 1C and D). After treatment of cells with centrinone B, p53 and p21 failed to accumulate in *TRIM37*^−/-^ cells at 200 or 500 nM centrinone B (Figure 3B). In the presence of supernumerary centrosomes, but not in the absence of centrosomes, MDM2 is cleaved via the ANKRD26/PIDDosome pathway that relieves its p53 inhibitory function and promotes p53 transcriptional activity [31, 32, 62]. We also noted a small amount of cleaved MDM2 [62] in untreated *TRIM37*^−/-^ cells with no additional increase after 200 nM centrinone B. This cleavage product was lost after treatment with 500 nM centrinone B. Thus, we find that TRIM37 does not affect gain or loss of centrosomes after centrinone B treatment but is required for the induction of both p53 and p21 in response to these treatments as previously observed [30], and is required for MDM2 cleavage.

**Figure 3.**
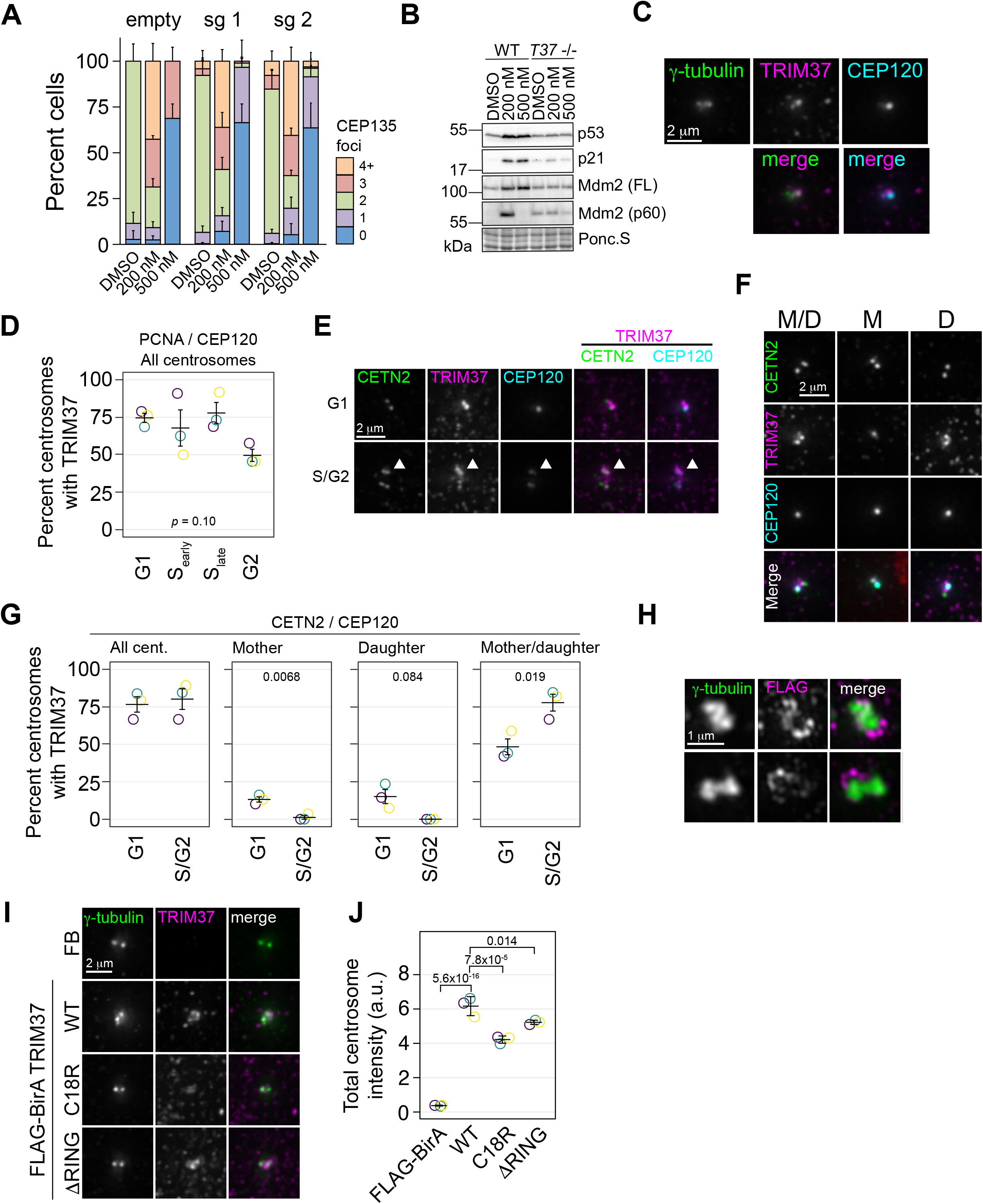
TRIM37 is a centrosome-associated protein. **(A)** RPE-1 Cas9 cells were stably infected with virus directing the expression of one of two sgRNAs against TRIM37 or empty vector. Selected cells were treated with DMSO, or 200 nM or 500 nM centrinone B for 4 days, fixed and stained for CEP135 and foci counted. (n = 3, N ≥169). **(B)** Cells from (A) were also processed for Western blotting using the indicated antibodies. FL – full length. p60 – p60 fragment. Ponc.S indicates total protein. **(C)** Asynchronous RPE-1 cells were fixed and stained with the indicated antibodies. Pairwise merged images are shown (bottom). **(D)** Asynchronus RPE-1 Cas9 cells were fixed and stained for TRIM37, PCNA, and CEP120. The number of TRIM37-positive centrosomes was manually determined for each cell cycle stage (n = 3, N ≥ 96). *p*-value from one-way Anova. **(E)** and **(F)** RPE-1 Cas9 cells were fixed and stained for the indicated antibodies. Examples of different cell cycles stages and TRIM37 localizations are shown in **(E)** and **(F)**, respectively. Arrowhead in **(E)** indicates TRIM37 preference for one of two centrosomes. M/D: Mother /daughter, M: Mother, D: Daughter **(G)** Quantification of cells shown in **(E)** and **(F)** (n = 3, N = ≥ 94). **(H)** RPE-1 *TRIM37*^−/-^ cells stably expressing FB-TRIM37 were fixed, stained with the indicated antibody and imaged using 3D-SIM. Two representative images are shown. **(I)** RPE-1 *TRIM37*^−/-^ cells stably expressing the indicated construct (FB = FLAG-BirA) were pre-extracted, fixed and stained for the indicated protein. **(J)** Centrosomal TRIM37 signal from cells in **(I)** was quantified. Means from each replicate are shown as open circles. Resulting mean and standard deviation shown (n = 3, N ≥ 84). Significant *p*-values from Dunnett post-hoc test using ‘WT’ as control after one-way ANOVA shown. Note that the results from (I) and (J) and those in Figure 7 – figure supplement 1C and E are from the same experiment therefore ‘FLAG-BirA’ and ‘WT’ are duplicated in these panels.

Since TRIM37 was a prey for multiple centrosome baits in our previous BioID survey of centrosomal proteins [63], we sought to determine if TRIM37 localized to centrosomes. Immunofluorescence staining of endogenous TRIM37 indicated that the protein was associated with the centrosome in most cells although it did not strictly co-localize with either γ-tubulin or CEP120 (Figure 3C). We verified that the anti-TRIM37 antibody used reliably detected endogenous TRIM37. The overall TRIM37 signal was greatly reduced in *TRIM37*^−/-^ cells (Figure 3 – figure supplement 1E) and TRIM37 detected at the centrosome in interphase cells was largely diminished and could not be reduced further using siRNA directed against TRIM37 (Figure 3 – figure supplement 3E-G). Since the intensity difference in mitotic cells was not as large that observed in interphase cells after disruption and/or knockdown, we restricted our analysis to interphase cells (Figure 3 – figure supplement 1E-G). To determine if TRIM37 localization was cell cycle-dependent, we co-stained TRIM37 with PCNA and CEP120 (Figure 3D) and detected TRIM37 in all cell cycle stages we could discern. We further determined if TRIM37 preferentially localized to mother or daughter centrioles by staining with CETN2 to detect all centrioles and CEP120 that preferentially localizes to daughter centrioles (Figure 3F and G). TRIM37 localized to both mother and daughter centrioles in most cells and localized exclusively to the mother or daughter in only a small percentage of cells. Interestingly, in cells with two centrosomes, TRIM37 appeared to favour one centrosome over the other (Figure 3E, arrowhead) and we observed a minor preference for exclusive association with mother or daughter centrioles in G1 cells (Figure 3G). Given that the fluorescence signal from detecting endogenous protein was too low for super-resolution imaging, we performed 3D-SIM on RPE-1 Cas9 *TRIM37*^−/-^ cells stably expressing FLAG-BirA-TRIM37 (FB-TRIM37) (Figure 3H, Figure 3 – figure supplement 2A-C). FB-TRIM37 formed partial ring structures preferentially surrounding one of the γ-tubulin foci. Moreover, the FB-TRIM37 signal was discontinuous with a dot-like distribution around the ring.

As a member of the TRIM family of E3 ligases, TRIM37 contains an N-terminal RING domain followed by a B-box and a coiled-coil region CCR [64]. The E3 ligase activity of TRIM37 has been implicated in its centrosomal-related functions in mitotic length [43], PCM stability [42, 43] and PLK4 localization [43]. We created two TRIM37 E3 ligase mutants, one containing a C18R point mutation [33] and another deleting the RING domain entirely (ι1RING). After stable expression in RPE-1 *TRIM37*^−/-^ cells, the steady state abundance of both ligase mutants was greater than the wild type (WT) protein, consistent with TRIM37 auto-regulating its stability [43] (Figure 3 – figure supplement 2A). Correspondingly, immunofluoresence for the tagged proteins confirmed their relative abundances and demonstrated that the proteins were expressed in all cells (Figure 3 – figure supplement 2B and C). The proteins were found primarily in the cytoplasm, sometimes in punctate structures possibly representing their peroxisomal localization [34]. Further, we quantified the centrosomal localization of these proteins and found that both E3 mutants localized to the centrosome, albeit at slightly lower levels than the WT protein (Figure 3I and J). Thus, TRIM37 localizes to an area near the centrosomes proper, surrounding the PCM in a manner that is partially dependent on its E3 ligase activity.

### An E3-independent TRIM37 activity mediates growth arrest after PLK4 inhibition

To determine if TRIM37 E3 ligase activity promotes growth arrest after centrinone B treatment, we performed clonogenic survival assays with the E3 mutant rescue lines in the presence of DMSO,200 nM or 500 nM centrinone B (Figure 4A). Expression of FB-TRIM37 fully restored the growth arrest triggered by centrinone B as did the expression of either C18R or ι1RING (Figure 4A). These data suggest that TRIM37 promotes growth arrest in response to centrinone B in an E3-independent manner.

**Figure 4.**
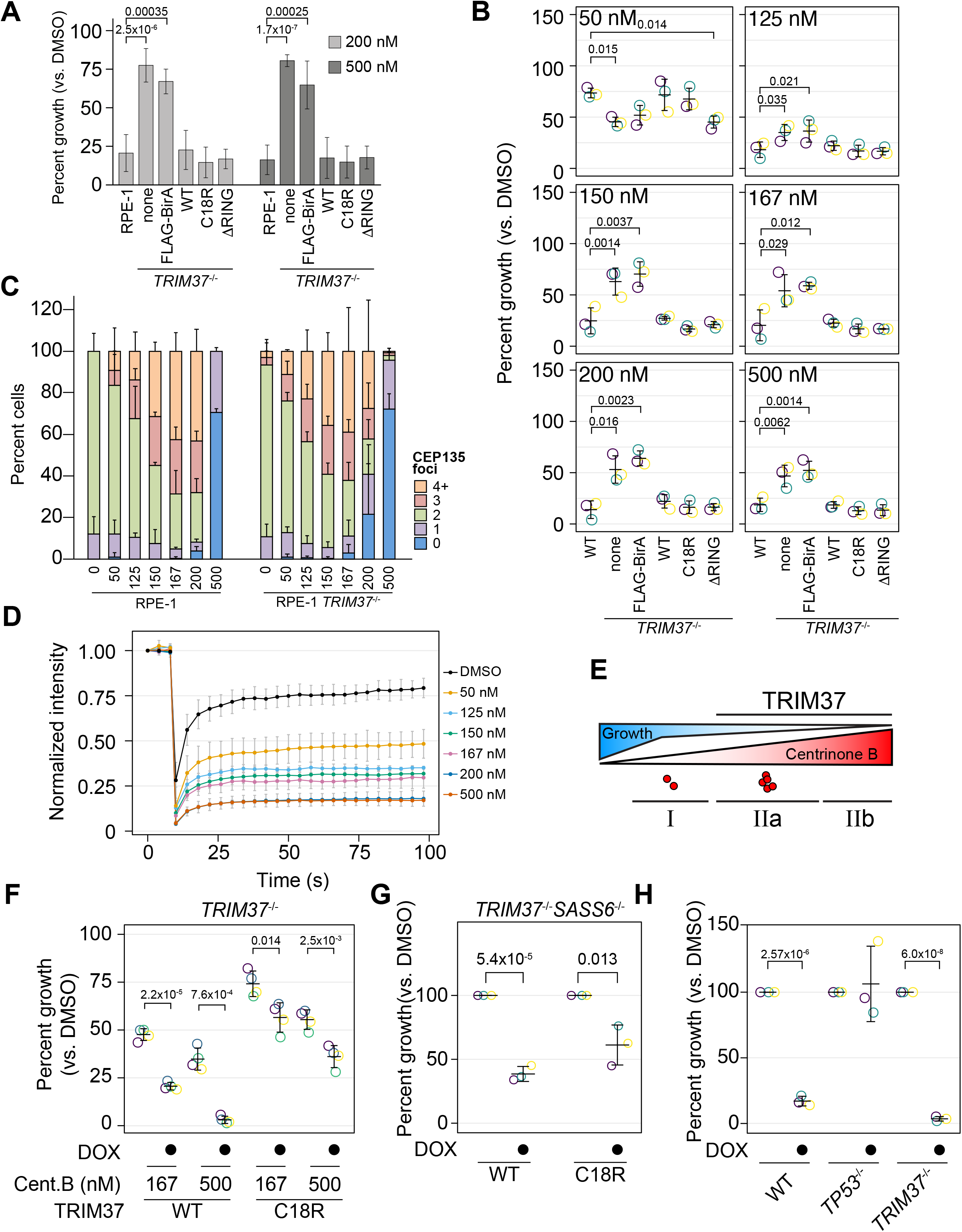
TRIM37 E3-independent activity is required for growth arrest. **(A)** WT RPE-1, *TRIM37*^−/-^ (none) and *TRIM37*^−/-^ cells expressing FLAG-BirA (FB) or the indicated FB-TRIM37 (WT, C18R, ΔRING) construct were seeded for clonogenic assays and grown in DMSO or the indicated concentration of centrinone B for 14 days. Colony density was quantified and growth compared to that in DMSO determined. Means and standard deviation shown (n = 3). **(B)** WT RPE-1, *TRIM37*^−/-^ (pool) and *TRIM37*^−/-^ expressing FB or the indicated FB-TRIM37 construct were seeded for clonogenic assays and grown in DMSO or the indicated concentration of centrinone B for 14 days. Colony density was quantified and growth compared to that in DMSO determined. Means from each replicate are shown as open circles. Resulting mean and standard deviation shown (n = 3). Significant *p*-values from Dunnett post-hoc test using ‘WT’ as control after one-way ANOVA shown. Note that the results from this experiment and those in Figure 7C are from the same experiment therefore ‘WT’, ‘*TRIM37*^−/-^ none’, ‘*TRIM37*^−/-^ FLAG-BirA’ and ‘*TRIM37*^−/-^ WT’ are duplicated in these panels. **(C)** WT or *TRIM37*^−/-^ (pools) RPE-1 cells were treated with DMSO (0) or the indicated concentration of centrinone B (nM) for 4 d before fixing and staining for CEP135. CEP135 foci per cell were manually counted. Mean and standard deviation shown (n = 3, N ≥ 55 per condition). **(D)** RPE-1 cells were transfected with GFP-PLK4kin+L1 and treated with DMSO or the indicated concentration of centrinone B for 16 h. The mean and standard deviation among the independent replicates is shown (n = 3, N ≥ 12). **(E)** Model showing growth inhibition ‘phases’. Growth is inhibited as a function of centrinone B. Phases dependent on TRIM37 are indicated. Red dots indicate centrosome number. **(F)** RPE-1 *TRIM37*^−/-^ cells expressing DOX-inducible TRIM37-3xFLAG or TRIM37 C18R-3xFLAG were seeded for clonogenic assays in the absence and presence of doxycycline and DMSO or the indicated concentration of centrinone B. After incubation for 14 d, colony density was quantified and growth compared to that in DMSO determined. Means from each replicate are shown as open circles. Resulting mean and standard deviation shown (n = 4). Significant *p*-values from pairwise t-tests comparing -DOX and +DOX samples are shown. **(G)** RPE-1 *TRIM37*^−/-^ *SASS6*^−/-^ cells expressing DOX-inducible TRIM37-3xFLAG or TRIM37 C18R-3xFLAG were seeded for clonogenic assays in the absence and presence of doxycycline. After incubation for 14 d, colony density was quantified and growth compared to that in DMSO determined. Means from each replicate are shown as open circles. Resulting mean and standard deviation shown (n = 3). Significant *p*-values from pairwise t-tests comparing -DOX and +DOX samples are shown. **(H)** The indicated RPE-1 line expressing inducible PLK4-3xFLAG were seeded for clonogenic assays in the absence and presence of doxycycline. After incubation for 14 d, colony density was quantified and growth compared to that in DMSO determined. Means from each replicate are shown as open circles. Resulting mean and standard deviation shown (n = 3). Significant *p*-values from pairwise t-tests comparing -DOX and +DOX samples are shown.

To corroborate these observations, we created *TRIM37* knockout pools in RPE-1 and A375 cells using an sgRNA distinct from that used to make the clonal line (Figure 3 – figure supplement 1C). These pools were infected with virus to express TRIM37 and the indicated TRIM37 mutants (Figure 4 – figure supplement 1A and B). In addition, we performed clonogenic assays using varying centrinone B concentrations to fully characterize the growth arrest activity promoted by TRIM37 (Figure 4B, Figure 4 – figure supplement 1C). Robust cell arrest of WT cells was observed after treatment with 125 nM centrinone B or greater. Correspondingly, in RPE-1 cells, we observed increases of p53 and p21 abundance with increasing centrinone B that was attenuated in *TRIM37*^−/-^ cells (Figure 4 – figure supplement 1D). Interestingly, cells lacking TRIM37 arrested after treatment with very low doses of centrinone B (50 - 125 nM) but only partially after higher concentrations (≥ 150 nM). As observed with cell lines derived from the clonal TRIM37 disruption, both E3-defective mutants, C18R and ΔRING, promoted growth arrest activity. To examine PLK4 function more closely after inhibition by centrinone B we monitored centrosome number and cellular PLK4 mobility. There was a dose-dependent increase in centrosome number up to 167 or 200 nM after which cells harboured one or no centrosomes (Figure 4C and Figure 4 – figure supplement 1E). Surprisingly, abnormal centrosome number did not correlate with robust growth arrest in WT cells; cell growth was almost completely inhibited at 125 nM centrinone B although we observed minor centrosome abnormalities at this concentration. Recently, PLK4 phosphorylation status near its phosphodegron sequence was linked to its cellular mobility where decreased phosphorylation at this site resulted in decreased mobility [65]. We expressed a PLK4 reporter construct (GFP-PLK kin+L1) and monitored PLK4 mobility using FRAP after treatment with centrinone B (Figure 4D) [65]. PLK4 mobility decreased with increasing centrinone B concentrations that mirrored growth arrest activity. Based on RPE-1 cells, our data uncovered three phases in the response to PLK4 inhibition. Phase I is TRIM37-independent and occurs at centrinone B concentrations where cells display minor centrosome number aberration (≤ 125 nM); a TRIM37-dependent phase II occurs at ≥ 150 nM centrinone B and can be further separated based on centrosome number; phase IIa cells harbour overduplicated centrosomes (150 – 200 nM) while phase IIb cells have lost one or both centrosomes (500 nM) (Figure 4E). Similar trends were observed using A375 cells, although the exact concentrations of centrinone B required differed between the cell lines (Figure 4 – figure supplement 1C and E).

We noted that our FB-TRIM37 construct was overexpressed compared to the endogenous protein (Figure 4 – figure supplement 2A), so we created *TRIM37*^−/-^ cells lines that expressed inducible TRIM37-3xFLAG or TRIM37C18R-3xFLAG. Compared to our stable cell lines, the inducible lines expressed TRIM37 closer to endogenous levels, although the C18R mutant was always more highly expressed, similar to previous studies (Figure 4 – figure supplement 2A - E) [43]. While the total cellular amount of the stable FLAG-BirA constructs was ∼20-fold higher than that of the DOX-inducible system (Figure 4 – figure supplement 2B), the centrosomal difference was only ∼ 3-fold (Figure 4 – figure supplement 2C). We performed clonogenic growth assays in the absence or presence of DOX and 0, 167 or 500 nM centrinone B (Figure 4F). Expression of either WT or C18R TRIM37 resulted in growth arrest although the growth arrest phenotype caused by TRIM37 C18R expression was weaker than that of WT cells. These data suggest that TRIM37 can support growth arrest after PLK4 inhibition in the absence of E3 ligase activity.

To distinguish between PLK4 activity and centrosome number, we used orthologous methods to control centrosome number. First, we disrupted the gene encoding SASS6, a protein required for centriole duplication [66, 67], to induce centrosome loss in *TRIM37*^−/-^ cells (Figure 4 – figure supplement 3A). We recovered cells lacking centrioles indicating that TRIM37 is also required for growth arrest in these conditions since *SASS6* is essential in WT cells [66]. Interestingly, *TRIM37*^−/-^ *SASS6*^−/-^ cells were completely resistant to any dose of centrinone B, unlike *TRIM37*^−/-^ alone (Figure 4 –figure supplement 3B). Similar to centrosome loss induced by PLK4 inhibition, inducible expression of TRIM37 C18R partially rescued the growth arrest phenotype caused by SASS6 loss (Figure 4G, Figure 4 – figure supplement 3C-D). To induce centrosome amplification, we overexpressed PLK4 (Figure 4 – figure supplement 3E). Similar to previous studies [31], we found that disruption of *TRIM37* was unable to suppress growth arrest in these conditions (Figure 4H). Together, these data suggest that TRIM37 is required for growth arrest after moderate or full PLK4 inhibition and this activity is partially independent of TRIM37’s E3-ligase activity.

### Growth arrest upon treatment with centrinone B does not correlate with an increase in mitotic length

Abnormal mitotic length has been put forth as an attractive model to explain growth arrest after centrosome loss [22]. We measured mitotic length after inhibiting PLK4 at different centrinone B concentrations using live-cell imaging of WT or *TRIM37*^−/-^ RPE-1 cells. Cells were treated with the indicated concentration of centrinone B for 3 days before imaging them for 24 h, and the length of mitosis from nuclear envelope breakdown (NEBD) to telophase was quantified (Figure 5A). Wild type RPE-1 cells treated with centrinone B did not exhibit a significant increase in mitotic length, compared to untreated cells, until concentrations reached 150 nM. Strikingly, the mitotic length of *TRIM37*^−/-^ cells treated with centrinone B was similar to that of WT cells until concentrations of 200 and 500 nM centrinone B; at which point the absence of TRIM37 partially suppressed the increase in mitotic length in WT cells [43]. We next examined whether TRIM37 E3 activity was required to rescue the shortened mitotic length observed in *TRIM37*^−/-^ cells. We induced the expression of WT TRIM37 and the C18R mutant in *TRIM37*^−/-^ cells, treated them with 500 nM centrinone B and measured mitotic length. In the absence of protein induction, we observed a slight increase in mitotic length after centrinone B treatment and a larger increase after induction of WT TRIM37 (Figure 5B). In contrast, the mitotic length in the presence of centrinone B after expression of TRIM37 C18R did not change compared to similarly treated uninduced cells. Thus, the increase in mitotic length observed after full PLK4 inhibition (500 nM centrinone B) appears to be dependent on TRIM37 E3 activity.

**Figure 5.**
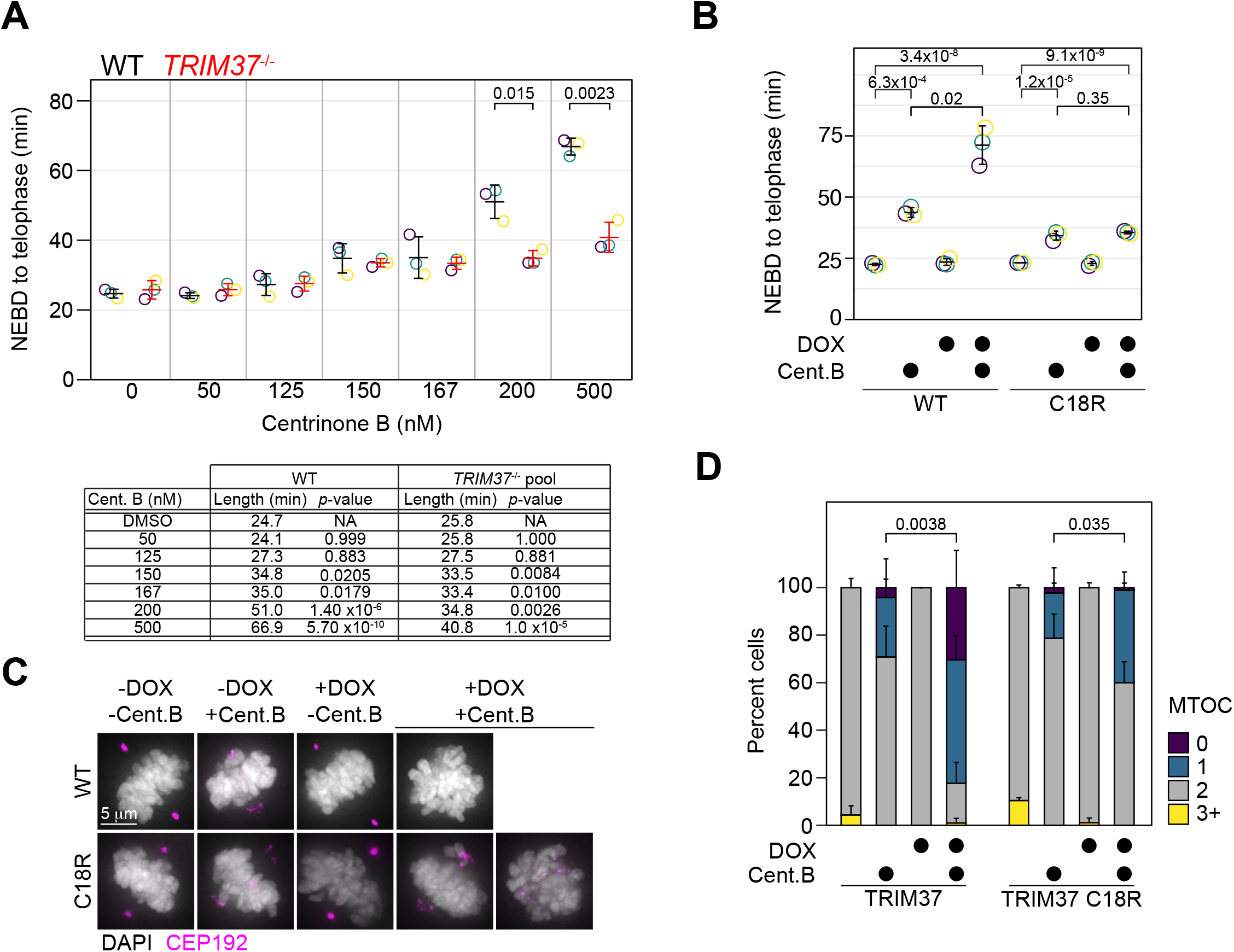
TRIM37 displays E3-dependent and -independent mitotic activities. **(A)** WT RPE-1 and *TRIM37*^−/-^ cells were incubated with DMSO (0) or the indicated concentration of centrinone B for 3 d before live imaging for 24 h. The time from nuclear envelope breakdown (NEBD) to telophase was determined. Means from each replicate are shown as open circles. Resulting mean and standard deviation shown (n = 3). Significant *p*-values from pairwise t-tests comparing WT and *TRIM37*^−/-^ samples are shown. (n = 3, N ≥ 30). Table below indicates average mitotic length and *p*-value from a Dunnett post-hoc test using ‘DMSO’ as control after one-way ANOVA. **(B)** RPE-1 *TRIM37*^−/-^ cells were treated with DMSO or 500 nM centrinone B (Cent.B) in the absence of presence of doxycycline (DOX) for 3 d before live imaging for 24 h. The time from nuclear envelope breakdown (NEBD) to telophase was determined. Means from each replicate are shown as open circles. Resulting mean and standard deviation shown (n = 3). Significant *p*-values from Dunnett post-hoc test using ‘uninduced, DMSO treated’ cells as a control after a one-way ANOVA are shown. (n = 3, N ≥ 46). **(C)** RPE-1 *TRIM37*^−/-^ cells expressing the indicated TRIM37 protein (WT or C18R) were treated with DMSO or 500 nM centrinone B (Cent.B) in the absence of presence of doxycycline (DOX) for 3 d before fixing and staining for CEP192. Representative images shown. **(D)** The number of discernable MTOCs characterized by the accumulation of CEP192 in cells from (C) was quantified. Means and standard deviation shown. (n = 3, N ≥ 29). For each TRIM37 construct the number of cells incubated with centrinone B and with 2 MTOCs in uninduced and induced samples was compared using a pairwise t-test.

In the absence of centrosomes, amorphous collections of primarily PCM components act as pseudo-MTOCs in cells lacking TRIM37 [30]. We induced the expression of WT TRIM37 or TRIM37 C18R in *TRIM37*^−/-^ cells, treated them with 500 nM centrinone B for 3 days before staining for CEP192 (Figure 5C) and analyzed mitotic cells for the number of MTOCs in each cell based on CEP192 distribution (Figure 5D). In DMSO-treated cells, all lines mostly formed two distinct MTOCs. In the presence of centrinone B, but in the absence of induced protein, most cells formed two fragmented MTOCs, characteristic of *TRIM37*^−/-^ cells after PLK4 inhibition [30]. Cells expressing WT TRIM37 and treated with centrinone B harboured a single fragmented MTOC or none at all. Lastly, cells expressing TRIM37 C18R and treated with centrinone B displayed a partial phenotype where some cells formed two dispersed MTOCs while others displayed a single dispersed MTOC. Thus, TRIM37 E3 ligase activity is not strictly required to suppress pseudo-MTOCs or to promote cell growth after centrosome loss.

### TRIM37 affects the abundance and localization of both centriolar and PCM components in an E3-dependent manner

To assess whether TRIM37 regulates the abundance of centriolar or PCM components, we initially probed for PCM components in RPE-1 and RPE-1 *TRIM37*^−/-^ lines but did not detect any significant changes in steady-state protein levels (Figure 6A and Figure 6 – figure supplement 1A). To determine if overexpressed TRIM37 affected steady state PCM levels, we again probed for PCM proteins in *TRIM37*^−/-^ cells stably expressing FLAG-BirA, FB-TRIM37 or the TRIM37 E3 mutants. We observed a significant decrease in CEP192 that was E3-dependent but did not observe significant changes in PCNT or CEP215 (Figure 6B and Figure 6 – figure supplement 1B). We next expressed inducible WT or C18R TRIM37 in WT RPE-1 cells [42, 43]. After expression of WT TRIM37 for 4 or 8 h we detected a 50 to 75% decrease in total CEP192 protein (Figure 6C, left panels). In all cases the decrease in CEP192 was E3-dependent. These results are consistent with previous observations that also indicated that TRIM37 is a negative regulator of CEP192 abundance [42, 43]. We extended these findings by using immunofluorescence to specifically detect changes at mitotic centrosomes (Figure 6 – figure supplement 1C). Quantification revealed that acute expression of WT TRIM37, but not C18R decreased the intensity of CEP192 and PCNT at the centrosome (Figure 6D, Figure 6 – figure supplement 6D). Interestingly the centriole component CEP120 was similarly diminished at the centrosome indicating that this effect is not specific to PCM proteins (Figure 6D). We find that over-expression of TRIM37 affects the overall abundance of CEP192 and the mitotic accumulation of CEP192, PCNT and CEP120 in an E3-dependent manner.

**Figure 6.**
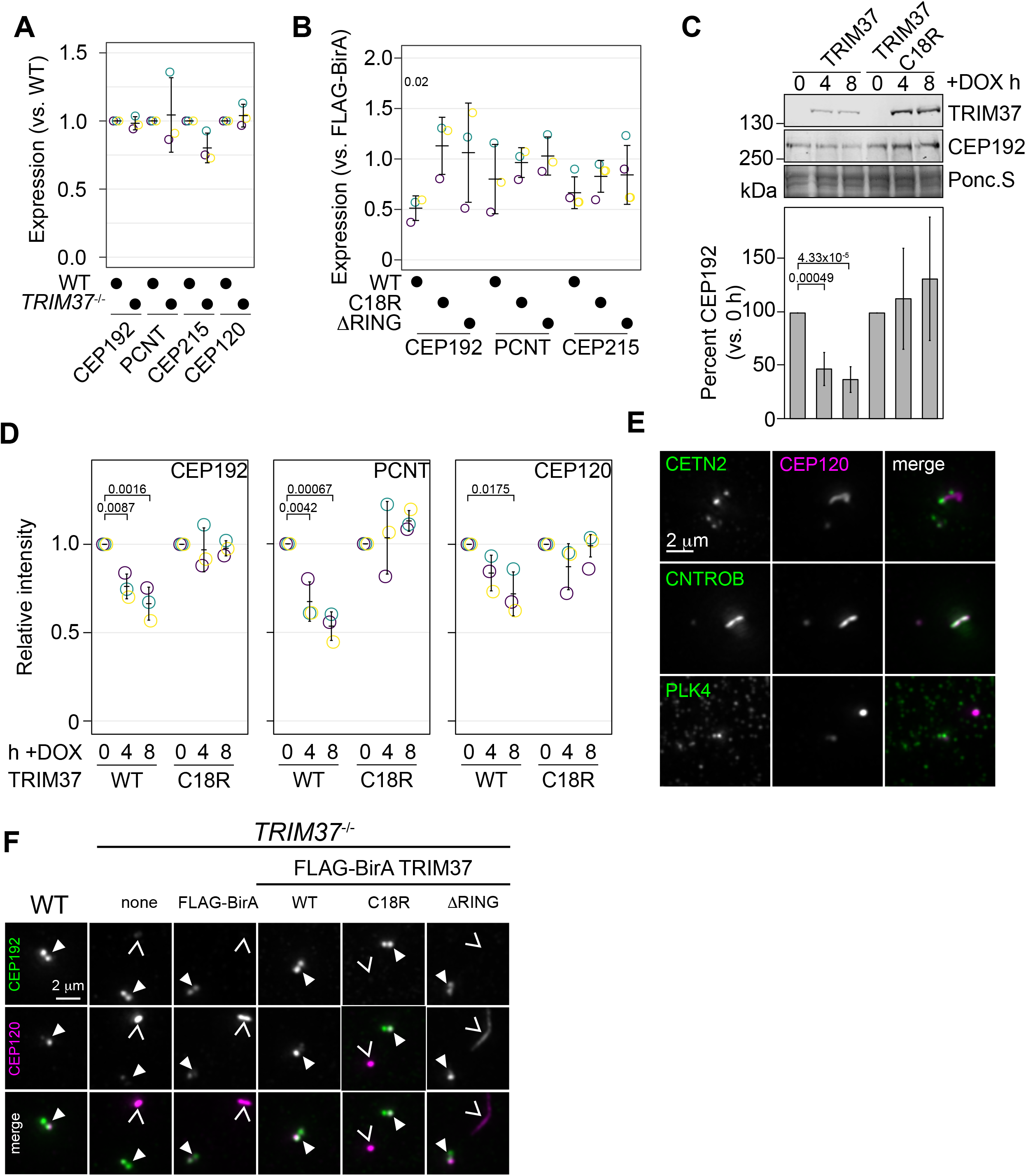
TRIM37 negatively regulates centriole and centrosome proteins. **(A)** WT RPE-1 and *TRIM37*^−/-^ cells were processed for Western blot and probed for the indicated proteins. Band intensity was quantified and expressed as expression compared to WT cells. Relative intensity from each replicate are shown as open circles. Resulting mean and standard deviation shown (n = 3). Data was tested by pairwise t-test between WT and *TRIM37*^−/-^. No significant differences were observed. **(B)** *TRIM37*^−/-^ RPE-1 cells stably expressing FLAG-BirA or the indicated FB-TRIM37 protein were processed for Western blot and probed for the indicated proteins. Band intensity was quantified and expressed as expression compared to cells expressing FLAG-BirA. Relative intensity from each replicate are shown as open circles. Resulting mean and standard deviation shown (n = 3). Significant *p*-values from Dunnett post-hoc test using FLAG-BirA as a control after a one-way ANOVA are shown. The band intensity from FLAG-BirA cells was set to ‘1’ and is omitted from the plot for clarity. **(C)** WT RPE-1 cells expressing doxycline-inducible TRIM37-3xFLAG (WT) or TRIM37 C18R-3xFLAG (C18R) were induced with doxycycline for 0, 4 or 8 h. At each time point, extracts were prepared and analyzed by Western blot for the indicated protein (right). Ponc.S indicates equal loading. CEP192 abundance was quantified and normalized to the intensity at time 0 h (bottom) Mean and standard deviation shown (n = 3). Significant *p*-values from Dunnett post-hoc test using time 0 h as a control after a one-way ANOVA are shown. **(D)** Cells from (C) were also fixed and immunostained for the indicated proteins. The centrosomal intensity from mitotic cells was determined. Intensity values were normalized to 0 h. Means from each replicate are shown as open circles. Resulting mean and standard deviation shown (n = 3, N = 60). Significant *p*-values from Dunnett post-hoc test using time 0 h as a control after a one-way ANOVA are shown. **(E)** RPE-1 *TRIM37*^−/-^ cells were fixed and stained for CEP120 and the indicated proteins. **(F)** RPE-1 (WT) or *TRIM37*^−/-^ cells stably expressing FLAG-BirA or the indicated FB-TRIM37 protein were fixed and stained for the indicated protein. Arrowhead indicates centrosome defined by CEP192. Caret mark indicates ectoptic structure defined by CEP120.

In the absence of TRIM37, some centriolar proteins form CTNROB-dependent ectopic intracellular aggregates, termed Cenpas, in interphase cells [40, 43]. We initially observed that CEP120 was mislocalized in *TRIM37*^−/-^ cells and co-localized with CNTROB (Figure 6E). Ectopic CNTROB structures remained after CEP120 depletion using siRNA suggesting that CEP120 is assembled downstream of CNTROB. (Figure 6 – figure supplement 2A). We found that CEP120 and CNTROB were detected in these structures and CETN2 foci accumulated near them (Figure 6E). Notably, we did not detect PLK4 in the CNTROB/CEP120 structure. Previously, PLK4 has been observed in these structures but could only be detected using a single antibody and the signal remained after siRNA treatment [30, 40, 41]. To determine if PLK4 could be recruited into these structures, we expressed PLK4-3xFLAG from an inducible promoter in *TRIM37*^−/-^ cells. PLK4-3xFLAG was not detected at non-centrosomal aggregates after 3 or 6 h induction using either anti-FLAG or anti-PLK4 antibodies, despite its accumulation at the centrosome (Figure 6 – figure supplement 2B). Additionally, these assemblies were not affected by the loss of PLK4 activity as they were observed in *TRIM37*^−/-^ cells treated with 500 nM centrinone B for 3 days (Figure 6 – figure supplement 2C and D). After stable expression of TRIM37 mutants in TRIM37^−/-^ cells these structures disappeared in an E3-dependent manner (Figure 6F, Figure 6 – figure supplement 2E). We find that CEP120 is a downstream component of the CNTROB structures formed in *TRIM37*^−/-^ cells and that the suppression of their formation requires TRIM37 E3 ligase activity.

### TRIM37 promotes the phosphorylation of PLK4

TRIM37 is suggested to associate with and ubiquitinate PLK4 [43]. We performed a structure-function analysis of TRIM37 to determine which region(s) of TRIM37 were required for PLK4 complex formation (Figure 7A) We transiently expressed a series of FB-TRIM37 deletion mutants and Myc-PLK4 in RPE-1 cells, immunoprecipitated the FB-TRIM37 constructs and probed for PLK4 (Figure 7A and B, Figure 7 – figure supplement 1A). Our results confirmed that PLK4 and TRIM37 form a complex in RPE-1 cells [43]. Further, the region from amino acids 505-709 of TRIM37 was sufficient to immunoprecipitate PLK4. Conversely, a TRIM37 mutant lacking this region (FB-Δ505-709) failed to pull-down PLK4. When stably expressed in *TRIM37*^−/-^ cells, FB-505-709 and FB-Δ505-709 were well expressed (Figure 7 – figure supplement 1B and D) but only FB-Δ505-709 localized to centrosomes (Figure 7 – figure supplement 1C and E). To determine if the PLK4/TRIM37 association was required for growth arrest activity we performed clonogenic assays using *TRIM37*^−/-^ cell lines stably expressing FB-TRIM37 505-709 and FB-TRIM37 1′505-709 (Figure 7C, Figure 7 – figure supplement 1B-E) or expressing inducible FB-1′505-709 (Figure 7 – figure supplement 1F and G). Surprisingly, the association between PLK4 and TRIM37 did not appear to be required for centrinone B-induced growth arrest.

**Figure 7.**
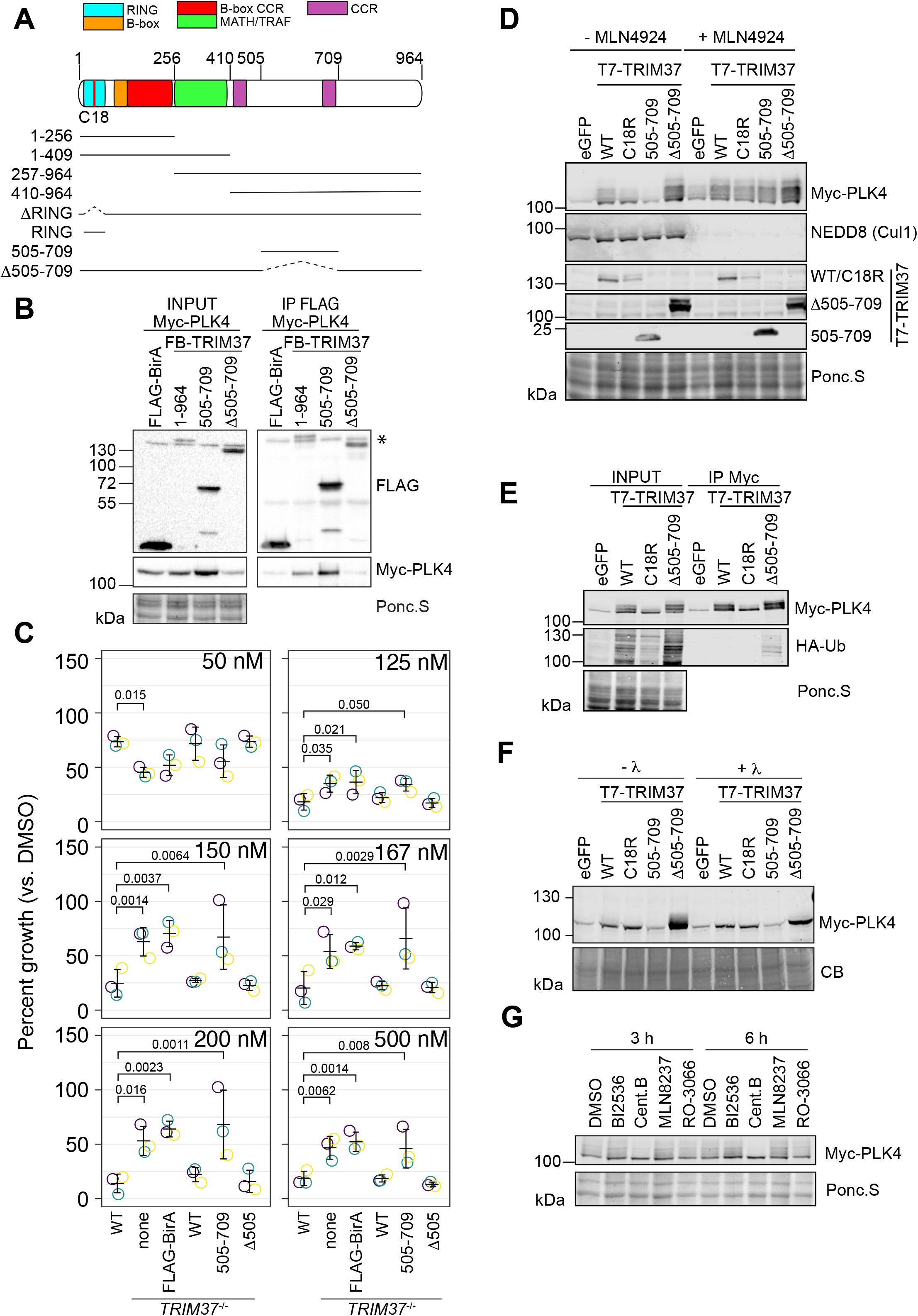
TRIM37 promotes PLK4 phosphorylation in an E3-dependent manner. **(A)** TRIM37 domain schematic. Constructs used for structure-function experiments indicated below. **(B)** RPE-1cells were transfected to express Myc-PLK4 and FLAG-BirA or the indicated FB-TRIM37 fusion protein (top). Cells were lysed and subjected to anti-FLAG immunoprecipitation. Input and immunoprecipitates were analyzed by immunoblotting for the FLAG-BirA fusions (FLAG) or for Myc-PLK4. Ponc.S indicates total protein. * indicates position of FLAG-Cas9. **(C)** WT RPE-1, *TRIM37*^−/-^ and *TRIM37*^−/-^ expressing FB or the indicated FB-TRIM37 construct were seeded for clonogenic assays and grown in DMSO or the indicated concentration of centrinone B for 14 days. Colony density was quantified and growth compared to that in DMSO determined. Means from each replicate are shown as open circles. Resulting mean and standard deviation shown (n = 3). Significant *p*-values from Dunnett post-hoc test using ‘WT’ as control after one-way ANOVA shown. Note that the results from this experiment and those in Figure 4B are from the same experiment therefore ‘WT’, ‘*TRIM37*^−/-^ none’, ‘*TRIM37*^−/-^ FLAG-BirA’ and ‘*TRIM37*^−/-^ WT’ are duplicated in these panels. **(D)** HEK293T cells transfected to express Myc-PLK4 and the indicated protein (top) were grown overnight and subsequently treated with DMSO or MLN4924 for 22 h and MG132 for the final 4 h. Cell extracts were prepared and probed by Western blot using the indicated antibodies. Ponc.S indicates total protein. **(E)** HEK293T cells were transfected to express Myc-PLK4, HA-Ub and the indicated protein (top). Cells were harvested after 48 h and subjected to immunoprecipitation using anti-Myc antibodies. Input and immunoprecipitates were analyzed by immunoblotting for PLK4 and HA-Ub. Ponc.S indicates total protein. **(F)** HEK293T cells were transfected to express Myc-PLK4 and eGFP or the indicated T7-TRIM37 protein (top) for 48 h. MG132 was added for the final 4 h. Lysates were mock treated (-λ)or incubated with λ-phosphatase (+λ) and subsequently subjected to immunoblot for PLK4. CB indicates total protein. **(G)** HEK293T cells were transfected to express Myc-PLK4 and T7-TRIM37 Δ505-709 for 48 h. Cells were treated with the indicated inhibitor (top) for 3 or 6 h and analyzed by immunoblot for PLK4. Ponc.S indicates total protein.

PLK4 protein abundance is tightly controlled by multiple post-translational modifications including phosphorylation and ubiquitination [14-16,65]. The co-expression of Myc-PLK4 and T7-TRIM37 in HEK293T cells resulted in modification of PLK4 (Figure 7D). The modification was partially dependent on TRIM37 E3 activity, was not observed when TRIM37 505-709 was expressed, and increased in the presence of TRIM37 1′505-709. MLN4924 is a general inhibitor of cullin-RING E3 ligases and treating cells with this compound should inhibit ubiquitination of PLK4 by SCF^β-TrCP^ [68]. Treatment of cells with MLN4924 resulted in stabilization of PLK4 but the modified forms remained (Figure 7D). To directly test if the observed modification was ubiquitinated PLK4, we co-expressed Myc-PLK4, T7-TRIM37 and HA-Ub in 293T cells. After immunoprecipitating PLK4 we probed for HA-Ub to detect ubiquitinated species (Figure 7E). Although we detected an E3-dependent increase in total ubiquitinated proteins in the input of cells expressing WT and Δ505-709 T7-TRIM37, we only detected low levels of HA-Ub conjugates in the anti-Myc immunoprecipitates, suggesting that PLK4 modification upon expression of TRIM37 may not be due to its ubiquitination. As an alternative possibility, we tested if the modified PLK4 bands were due to phosphorylation by treating cell lysates with 11-phosphatase (Figure 7F). The slower migrating forms of PLK4 were lost after phosphatase treatment indicating that these modifications are primarily due to phosphorylation. To identify the kinase(s) responsible for the modification we treated cells with inhibitors targeting PLK4, AURKA, PLK1 and CDK1 for 3 to 6 h and probed for PLK4. The only treatment that substantially reduced PLK4 phosphorylation was PLK4 inhibition (Figure 7G). Since we observed that TRIM37 promotes PLK4 phosphorylation (Figure 7G) we monitored GFP-PLK4 mobility by FRAP in WT, *TRIM37*^−/-^ cells and after TRIM37 siRNA but did not observe any differences compared to control cells indicating that the phospho-forms of PLK4 stabilized by TRIM37 likely lie outside the phosphodegron region (Figure 7 – figure supplement 1H and I). Together these data suggest that TRIM37 promotes the accumulation of phosphorylated PLK4 in an E3-dependent manner but this phenomenon does not require robust interaction with PLK4 itself.

## DISCUSSION

Most animal cells harbouring abnormal centrosome numbers are subject to p53-dependent growth arrest and the mechanisms of these pathways are beginning to be understood. Here, by leveraging the various phenotypes caused by treatment of cells with different concentrations of the selective PLK4 inhibitor centrinone B, we uncover multiple pathways leading to growth arrest in response to abnormal centrosome numbers. Not only does centrosome number play a role, but we hypothesize that properties or the activity of PLK4 itself can also trigger growth arrest. Curiously, we found that TRIM37 is required for growth arrest in some, but not all centrinone B concentrations tested.

Several observations support our hypothesis that differential PLK4 inhibition used for our screens resulted in distinct cellular states. First, we observed clear differences in centrosome number. Second, MDM2 was cleaved using 200 nM, but not 500 nM centrinone B consistent with excess centrosomes in the former condition. Last, the genes derived from each screen were distinct. Specifically, we identified ANKRD26/PIDDosome only in the presence of excess centrosomes and also identified centriole proteins that, when disrupted, could decrease centrosome load, although we did not formally test this. We note that 200 nM centrinone B was not optimal to induce excess centrosomes in A375 cells (compare Figure 4C with Figure 4 – figure supplement 1E), yet we still identified genes that overlapped with those from the comparable RPE-1 screen suggesting that these cells were subjected to similar conditions.

Centrosome amplification after partial inhibition of PLK4 has been previously observed using CFI-400495 [45], YLT-11[69] or analog-sensitive alleles of *PLK4* [6]. Current models suggest that partially inhibited PLK4 reduces its auto-phosophorylation required for its degradation. As a consequence, PLK4 accumulates and promotes centriole overduplication [70]. While TRIM37 is involved in mediating growth arrest to partial PLK4 inhibition, it is not required for arrest after PLK4 over-expression, which also leads to extra centrosomes [31]. The fundamental difference between centrosome amplification caused by PLK4 over-expression or by partial inhibition may relate to the per molecule activity of PLK4, perhaps pointing a role for TRIM37 in regulating PLK4 activity. Our FRAP data indicated a dose-dependent decrease in phosphorylated, and therefore active, PLK4 upon increasing centrinone B that correlated well with growth arrest activity. If altered PLK4 activity, and not extra centrosomes, is responsible for growth arrest, why would we identify proteins such as ANKRD26 and the PIDDosome that have clear roles in response to supernumerary centrosomes [31, 32, 62]? Growth suppression after PLK4 inhibition at any concentration of centrinone B was only partially TRIM37-dependent, suggesting that multiple pathways might be activated in these conditions, one dependent on centrosomes and the other dependent on PLK4. Comparing our dataset with that of the PLK4 overexpression screen [31] yields an overlap of 22 genes that we propose are involved in a response to supernumerary centrosomes (Figure 2 – figure supplement 2G). We suggest that the genes unique to our dataset (i.e. low-dose centrinone B treatment) might modulate the response to inhibited PLK4. *CEP85* and *USP9X* are such genes and both encode proteins that affect STIL, a regulator of PLK4 activity. CEP85 is required for robust STIL interaction with PLK4 while USP9X stabilizes STIL [8, 71]. Interestingly, 53BP1 and USP28 were similarly dispensable for growth arrest after PLK4 overexpression [31]. In this case, the MDM2 p60 fragment that is known to interact with and stabilize p53 [72] might be redundant with the function of 53BP1-USP28. It is therefore an exciting possibility that PLK4 activity, by itself, may be a determinant of p53-dependent cell cycle arrest.

TRIM37 is proposed to mediate the degradation of CEP192 that in turn, affects PCM assembly in mitotic cells treated with PLK4 inhibitors [41, 42], however, there is no consensus on the exact centrosome or centriole TRIM37 targets. We observed a decrease in overall CEP192 protein levels, but not those of PCNT, CEP215 or CEP120 after stable expression of FB-TRIM37 (Figure 6B). Others have reported decreases in the protein levels of CEP192, but some did not detect decrease in the levels of CEP215 or CEP152 [41], whereas others have failed to detect changes in =CEP215 and PCNT protein abundance [42]. We did not detect reciprocal changes in protein levels in *TRIM37*^−/-^ cells (Figure 6A) suggesting that the effects on PCM are dependent on overexpressed TRIM37 or that the *TRIM37*^−/-^ cell line acquired a genetic or epigenetic change that suppresses effects on PCM proteins. TRIM37 overexpression also decreased the amount of CEP192, PCNT and CEP120 detected at mitotic centrosomes (Figure 6D and Figure 6 – figure supplement 1D). More work will be needed to identify direct and indirect targets of TRIM37 at the centrosome.

The E3 ligase activity of TRIM37 was required for changes in the bulk abundance of CEP192 and for the reduction in CEP192, PCNT and CEP120 proteins at mitotic centrosomes. In contrast, we observed dosage-dependent phenotypes for the E3 ligase mutant TRIM37 C18R. Stable and high expression of this variant caused a strong growth arrest phenotype (Figure 4A) while lower expression using an inducible system resulted in partial growth arrest activity in response to centrinone B treatment (Figure 4E and F). To explain this phenomenon, we consider that either the expression of the C18R mutant binds to and sequesters TRIM37 targets to phenocopy the effect of degradation, or that higher levels of TRIM37 C18R drive the formation of an E3-independent complex important for growth arrest, as has been suggested for a TRIM37 ligase-independent role in autophagy [38].

Current models of TRIM37 growth arrest function after PLK4 inhibition have primarily focused on mitotic length [22, 30, 42, 43]. After complete PLK4 inhibition, cells containing one or no centrosome arrested, but in these studies, mitotic length and growth arrest were not well correlated and not long enough to activate a ‘mitotic timer’ [23, 73]. This led to the speculation that small increases in mitotic length over multiple cell cycles might be equivalent to a single mitosis with a larger delay [22, 30] but this possibility has not yet been tested. Here however, we provide multiple lines of evidence that mitotic length may not be critical for growth arrest after PLK4 inhibition. First, in WT RPE-1 cells, we observed that treatment with 50 nM and 125 nM centrinone B led to ∼25% and 90% decreases in cell proliferation, respectively (Figure 4B) without causing concomitant increases in mitotic length (Figure 5A). Second, when comparing WT and *TRIM37*^−/-^ cells, we observed a significant difference in growth between these cell lines at 125 nM centrinone B or greater but did not detect significant differences in mitotic length until a treatment with 200 nM centrinone B. Third, induced expression of TRIM37 C18R caused a partial reduction in cell proliferation without any apparent effect on mitotic length (Figure 4E and F). We also note that PLK4 overexpression in mice or flies results in increased mitotic indices [25, 74] indicative of lengthened mitoses, yet TRIM37 is not required for growth arrest under similar conditions in cultured cells. Mitotic length after treatment with 200 nM and 500 nM centrinone B is clearly affected by TRIM37 but our data does not suggest that this directly influences growth arrest.

Although others have observed TRIM37-dependent ubiquitination of PLK4 [43], we detected low amounts of ubiquitinated PLK4 only after co-expression with the TRIM371′505-709 mutant (Figure 7E). In contrast, we found that TRIM37 primarily promoted PLK4 phosphorylation in a manner that was dependent on PLK4 activity itself (Figure 7F and G). We did not find that PLK4 mobility by FRAP was affected by the loss of TRIM37 in the absence or presence of centrinone B (Figure 7 – figure supplement H and I) suggesting that the phosphorylation sites stabilized by TRIM37 lie outside the PLK4 phosphodegron-adjacent regions monitored by this method [65]. It will be of interest to determine how TRIM37 promotes PLK4 phosphorylation, which regions of PLK4 are modified and to assess if these phosphorylation events are contributing to TRIM37-dependent growth arrest in response to abnormal centrosome numbers or altered PLK4 activity.

We described three distinct growth arrest phases after PLK4 inhibition characterized by cellular centrosome number abnormalities (Figure 4E). Our FRAP assays indicated that the mobility of PLK4 decreased after centrinone B treatment in a dose-dependent manner that mirrored the growth arrest activity of WT cells. We therefore propose that a direct aspect of PLK4 activity, either PLK4 itself or a substrate of PLK4, underlies the growth arrest after PLK4 inhibition (Figure 8). A level of PLK4 inhibition that does not affect centriole number initiates p53 arrest independent of TRIM37, but cell arrest after further PLK4 inhibition becomes dependent on TRIM37. TRIM37 itself promotes phosphorylation of PLK4 and can affect the abundance and/or localization of CEP192, PCNT and CEP120, but the effect of these functions are not clear. SASS6 was required for growth arrest at all centrinone B concentrations (Figure 4 – figure supplement 3B) suggesting that centrioles themselves may play a role to integrate the growth arrest signal.

**Figure 8.**
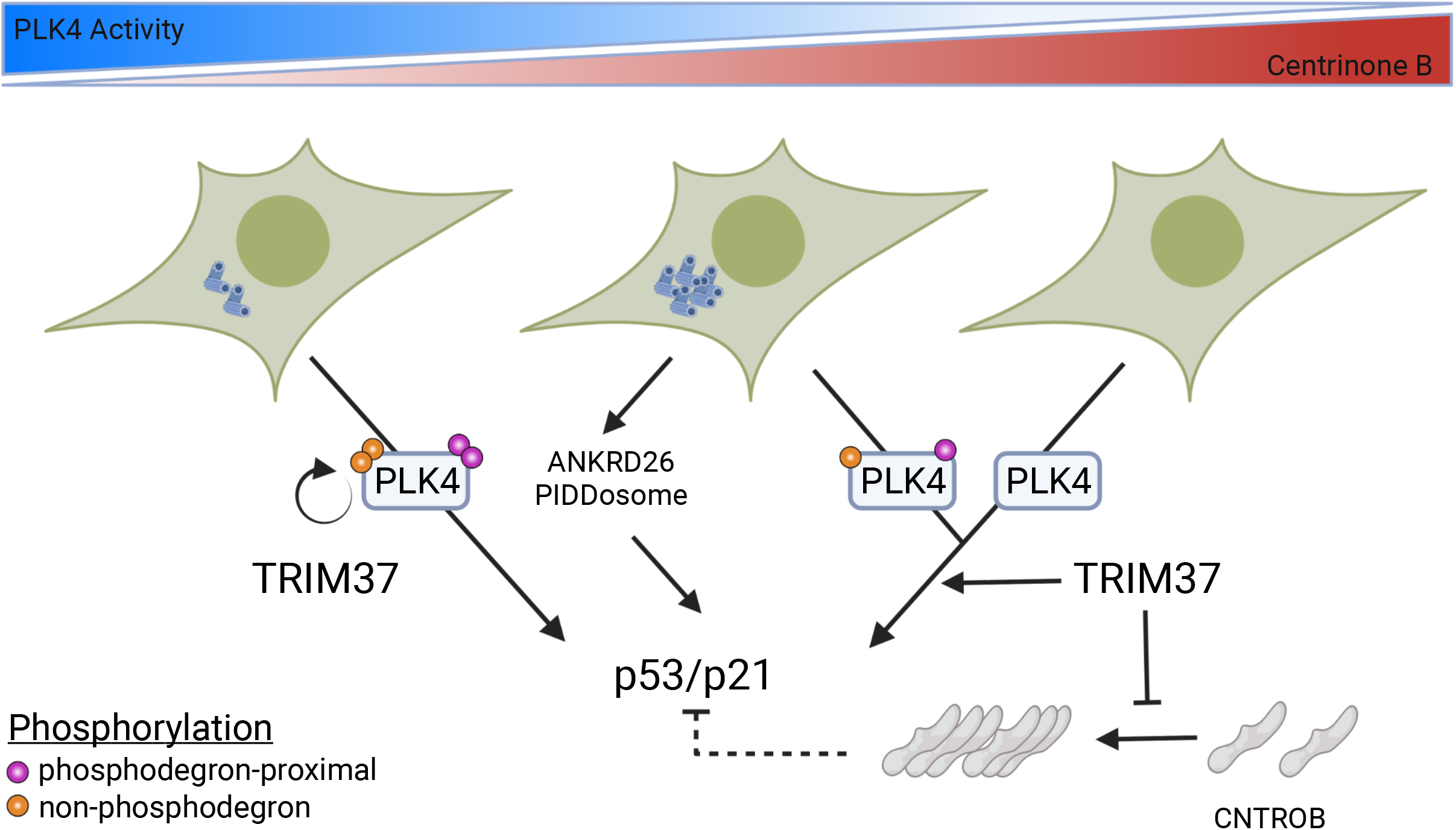
Model for growth arrest and TRIM37 growth arrest activity. PLK4 activity decreases in a dose-dependent manner upon centrinone B addition. TRIM37 promotes PLK4 auto-phosphorylation (orange circles) outside the phosphodegron region (purple circles). PLK4 inhibition initially results in TRIM37-independent growth arrest. Continued addition of centrinone B results in centrosome overduplication that is detected by the ANKRD26/PIDDosome pathway in addition to a TRIM37-dependent growth arrest pathway. Complete inhibition of PLK4 results in TRIM37-dependent growth arrest. TRIM37 also prevents the appearance of CNTROB-dependent aggregates. We hypothesize that these aggregates might affect p53/p21 activation (dotted lines) (Created with BioRender.com).

In closing, we used CRISPR/Cas9 screening to characterize the global, dose-dependent response to PLK4 inhibition. While previous studies focused on complete PLK4 inactivation and mitotic length after centrosome loss as a potential mechanism driving p53-dependent cell cycle arrest, we found that the loss of PLK4 activity better correlates with the subsequent growth arrest. Proteomic analysis of PLK4 substrates and the cellular aggregates that form in the absence of TRIM37 will be required to provide mechanistic details of this pathway and may yield the identification of PLK4 substrates underpinning this response.

## ACKNOWLEDGEMENTS

We thank members of the Pelletier lab for scientific feedback during the preparation of this work. We are grateful to Sally Cheung and Suzanna Prosser for proofreading the manuscript. The rabbit and rat CEP120 antibodies and CEP135 antibody were generous gifts from Li-Heui Tsai, Moe Mahjoub and Alex Bird, respectively. The TKOv1 virus library was generously provided by Jason Moffat. The pcDNA3 HA-ubiquitin plasmid was constructed by Abdallah Al-Hakim. pcDNA5 GFP-PLK4 and GFP-PLK4kin+L1 were gifts from Daiju Kitagawa. This work was funded through a Canadian Institute for Health Research Foundation Grant, an ORF RE-5 grant from the Ontario Ministry for Research, and a grant-in-aid from the Krembil Foundation. L.P. holds a Tier 1 Canada Research Chair in Centrosome Biogenesis and Function.

## AUTHOR CONTRIBUTIONS

L.P and J.M.T conceived the project and designed experiments. J.S and J.M.T. performed the CRISPR/Cas9 screens. J.M.T. performed all other experiments. R.P. optimized and performed FRAP experiments. A.S. performed FRAP experiments. L.P. and J.M.T. interpreted and analyzed the results. L.P. and J.M.T wrote the manuscript. L.P., J.M.T, R.P., A.S., J.S. and D.D. reviewed and edited the manuscript. D.D. supervised J.S.

## MATERIALS AND METHODS

### Cell culture and drug treatments

All cell lines were cultured in a 5% CO2 humidified atmosphere at 37°C. HEK293T (female, human embryonic kidney epithelial) cells are from ATCC. hTERT-RPE1 (female, human epithelial cells immortalized with hTERT) and A375 cells (female, human malignant melanoma epithelial) stably expressing Cas9 were a kind gift from D. Durocher. All references to RPE-1 and A375 cells herein refer to hTERT-RPE1 or A375 stably expressing Cas9. hTERT-RPE1, HEK293T and A375 were grown in Dulbecco’s Modified Eagle’s Medium (DMEM; Gibco) supplemented with 10% (v/v) fetal bovine serum (FBS; Gibco) and 2 mM Glutamax (Gibco). PLK4 inhibitor centrinone B (Tocris) was used as described. Nutlin-3a (Cayman Chemical) was used at 600 nM. The CDK1, PLK1 and Aurora A kinase inhibitors RO-3306 (Enzo Life Sciences), BI-2536 (ChemieTek) and MLN8237 (Selleck Chemicals) were used at 10 μM, 100 nM and 200 nM, respectively. MG132 (Selleck Chemicals) was used at 10 μM. G418 (WISENT Bioproducts) was used at 600 μg/mL for cell selection and 200 μg/mL for routine culture.

### Virus production

To produce lentivirus, 4×10^6^ HEK293T were seeded in a T-75 flask and subsequently transfected with 4 μg of the appropriate transfer vector, 3 μg psPAX2 and 2μg pCMV-VSV-G using 18 uL each Lipofectamine 3000/P3000 reagent (Invitrogen). After 24 h, growth medium was replaced with fresh medium containing 30% FBS and viral supernatant was collected after a further 48 h. Virus was stored at −80°C.

### CRISPR/Cas9 screening

CRISPR screens were performed as described [47, 75]. Briefly, Cas9-expressing cells were transduced with the TKOv1 viral library (∼90 k sgRNA) [47] at low MOI (∼0.3) in the presence of 4 μg/mL polybrene. RPE-1 cells were selected as described [76]. A375 cells were selected using 2 μg/mL puromycin. 10×10^6^ cells were harvested four days post-transduction and represents day 0. Cells were grown for 6 days before being split for drug treatment in technical triplicate and further grown for 21 days. A library coverage of > 100 cells/sgRNA was maintained at each step. gDNA from cell pellets was isolated using a QIAamp Blood Maxi Kit (Qiagen) and genome-integrated sgRNA sequences were amplified using the KAPA HiFi HotStart ReadyMix (Kapa Biosystems). Sequencing libraries were made by addition of i5 and i7 multiplexing barcodes in a second round of PCR and the product gel purified using QIAquick Gel Purification kit (Qiagen). Libraries were sequenced using Illumina HiSeq2500 or NextSeq500. Sequence data was analyzed using MAGeCK [49] to determine sgRNA distribution among the samples. Drug-treated samples at 21 days post-drug addition were compared to DMSO-treated cells at 12 days post-drug addition to equalize the number of cell doublings. Genes with FDR < 0.05 were used for further analysis. The significant gene list for the RPE-1 200 nM screen is the union from two independent biological replicates.

### Network analysis and gene enrichment

High-scoring genes from MAGeCK analysis were visualized using Cytoscape [77]. General node arrangement was performed using the yFiles Organic Layout and manually modified to facilitate visualization. Each screen condition (200 nM centrinone B, 500 nM centrinone B and Nutlin-3a) was considered as a source node, corresponding hits as target nodes and FDR as edge attributes. Genes from the indicated data sets were analyzed using the ClueGo app within Cytoscape [78]. Enrichments for Biological Function (circles) or Cellular Component (hexagons) based on all experimental evidence was determined. Only pathways with p-value < 0.05 are shown. Nodes arranged using the yFiles Organic Layout.

### CRISPR/Cas9 gene disruption

For lentivirus-mediated gene disruption of *TRIM37*, sgRNA sequences were cloned into plentiGuide-Puro as described [79]. RPE-1 Cas9 cells were infected with lentiviral particles and selected as described above for CRISPR/Cas9 screening. A Clonal *TRIM37*^−/-^ line was generated using *in vitro* transcribed (IVT) sgRNA. IVT templates were created by PCR using cr_tracrRNA, IVT forward, IVT reverse and sgRNA-specific oligonucleotides (TRIM37 sgRNA 1). PCR products were used directly as templates for IVT using HiScribe T7 transcription kit (NEB). Resulting RNA was purified using RNAClean XP beads (Beckman Coulter) and used to transfect RPE-1 cells using RNAiMAX (ThermoFisher) according to manufacturer’s instructions. Clonal lines were generated by limiting dilution and assessed for gene disruption by Western blot and TIDE [80] or Synthego ICE (Synthego Performance Analysis, ICE Analysis. 2019. v2.0. Synthego; accessed 9/19/2018) analyses. *TRIM37*^−/-^ pools in RPE-1 and A375 cells were generated similarly using an sgRNA targeting exon 5. After transfection, cells were grown in medium containing 500 nM centrinone B for two weeks to select for *TRIM37* disruption before growth in normal medium.

### Stable cell line generation

To generate cell lines, 200 000 cells were seeded with serial aliquots of viral supernatant and 4 μg/mL polybrene (MilliporeSigma) in one well of a 6-well plate. Medium was changed after 24 h and appropriate drug selection was added after an additional 24 h where required. For stable expression of FLAG-BirA rescue contstructs, immunofluorescence was performed to ensure all cells expressed the appropriate transgene. Doxycycline-inducible lines were selected with 600 μg/mL G418 until control cells died. We used pools that showed approximately 30% survival after initial selection.

### siRNA conditions

For siRNA knockdown experiments, 200 k cells were seeded per well of a 6-well plate. Cells were reverse transfected using the indicated siRNA trigger (Horizon Discovery, Dharmacon; Table S3). For each well, 5 μL of 20 μM siRNA was combined with 3 μL Lipofectamine RNAiMAX (ThermoFisher) in 125 uL OPTIMEM medium (Gibco). Media was replaced after 24 h and cells processed after 72 h.

### Immunofluorescence staining and microscopy

Cells were grown as indicated on No. 1.5 coverslips, washed once with PBS and fixed with −20°C methanol for at least 10 min. All subsequent steps performed at room temperature. Coverslips were rinsed with PBS and blocked with antibody solution (PBS, 0.5% (w/v) BSA and 0.05% Tween-20) for 15 to 30 min. Samples were incubated with primary antibodies (Table S3) for 1 h, washed 3 x 5 min and incubated with secondary antibodies (Table S3) and DAPI (0.1 μg/mL) for 45 min. Coverslips were washed 3 x 5 min and mounted on slides using Prolong Gold (Invitrogen). Deconvolution wide-field microscopy was performed using the DeltaVision Elite system equipped with an NA 1.42 60x PlanApo objective (Olympus) and an sCMOS 2048 x 2048 camera (Leica Microsystems). Each field was acquired with a z-step of 0.2 μm through the entire cell and deconvolved using softWoRx (v6.0, Leica Microsystems). Maximum intensity projections are shown (0.1080 μm/pixel). Display levels are the same for all images in a panel unless otherwise indicated.

For live imaging, 15 000 cells were seeded per well in an 8-well Lab-Tek II chamber slide. The next day fresh medium containing drug was added and cells incubated for 3 days. Fresh medium containing indicated drug and 200 nM SiR-DNA (Spirochrome) was added for 2 h before imaging. Microscopy was performed using the DeltaVision Elite system equipped with an NA 0.75 U Plan S-Apo objective (Olympus) and an sCMOS 2048 x 2048 camera. Each field was acquired with 6 x 2 μm z-step every 5 min for 24 h. The time between nuclear envelope breakdown and full chromosome separation judged by nuclear morphology was quantified.

Super-resolution microscopy was performed on a three-dimensional structured-illumination microscope (3D-SIM) (OMX Blaze v4, Leica Microsystems) as described [81].

### Image analysis

All automated quantification pipelines were created using CellProfiler 3.0 [82] (www.cellprofiler.org).

Figure 1A and B: Nuclei were detected using the DAPI channel and objects subsequently used as a mask to measure intensity in p53 or p21 channels. An arbitrary cut-off based on the distribution of p21 or p53 intensities in untreated cells was used to score positive cells.

Figure 3J, Figure 3 – figure supplement 1F and G, Figure 4 – figure supplement 2C: A centrosomal (γ-tubulin) or centriole marker (CEP120) were used to define centrosome regions. The TRIM37 images were masked by the centrosome objects and total intensity was measured. Figure 3 – figure supplement 2C and Figure 7 – figure supplement 1D: Nuclei were detected using the DAPI channel. The nuclear objects were expanded and a ring surrounding each nucleus was used as a mask to measure the total intensity in the BirA channel. The mean and standard deviation of the measured intensities of control cells was determined and a cut-off of the mean + 2.5x the standard deviation was used to score positive cells.

Figure 6D and Figure 6 – figure supplement 1D. Each image was manually cropped to include a single mitotic cell. Each channel was background subtracted using the lower quartile intensity of the entire image and each channel was segmented into objects using a robust background thresholding and the integrated intensity of each object was measured.

### Western blot

Cells were grown as indicated, washed once with PBS and resuspended directly in 2x SDS-PAGE sample buffer containing Benzonase (0.25 U/μL, MilliporeSigma) and heated at 95°C for 5 min. Proteins (typically 10 to 20 μg) were separated by SDS-PAGE and transferred to PVDF using a wet transfer apparatus (Bio-Rad). Total protein was detected by staining with PonceauS (MilliporeSigma) and scanning. All steps performed at room temperature unless indicated. Blocking and primary antibody incubations were performed using TBS-T (TBS + 0.05% Tween-20) with 0.5% skim-milk powder (Bioshop). Membranes were blocked for 30 min and incubated with primary antibody (Table S3) overnight at 4°C. After washing 3 x 5 min, membranes were incubated with secondary antibodies for 45. HRP-conjugated secondary antibodies (Bio-Rad) were incubated in TBS-T/milk for 45 min and washed 3 x 5 min with TBS-T before detecting using a Chemidoc imager (Bio-Rad). NearIR-conjugated secondary antibodies (LI-COR Biosciences) were incubated in TBS-T/milk + 0.015% SDS for 45, washed 3 x 5 min with TBS-T and 1 x 5 min with TBS before drying the membrane for 2 h at RT. Dried blots were imaged using an Odyssey CLx imager (LI-COR Biosciences).

Quantification of Western blots were performed on images obtained using NearIR secondary antibodies. Images were quantified using Image Studio software (LI-COR Biosciences) and normalized to Ponceau S or Coomassie Blue staining of the same lane.

### Clonogenic survival assays

Two-hundred and fifty RPE-1 or 200 A375 cells were seeded in either a 10 cm dish or 6-well plate. The next day medium was removed and medium containing the indicated drug was added. For experiments using doxycycline-inducible cell lines, the media was refreshed every 3 to 4 days to ensure continued expression of the induced proteins. After 12 to 14 days, plates were rinsed once with PBS and fixed and stained with 0.5% crystal violet (MilliporeSigma) in 20% methanol for at least 20 min. Plates were washed extensively with water, dried and scanned. Images were segmented using the Trainable Weka Segmentation tool [83] in ImageJ. A new model was built for each replicate if required. The resulting segmentation image was thresholded and used as a mask to overlay the original image that was inverted and background subtracted using a 50-pixel rolling circle, or the average of a region not containing colonies. The colony intensity per well or dish was then measured within the masked region.

### Immunoprecipitation and protein treatments

To detect complex formation between PLK4 and TRIM37, 2×10^6^ RPE-1 cells were seeded per 10 cm dish and transfected with 3.75 μg Myc-PLK4 and 3.75 μg pcDNA5 FLAG-BirA construct using 15 μL Lipofectamine 3000/P3000 (ThermoFisher). Cells were harvested 24 h post-transfection, washed once with PBS and resuspended in lysis buffer (50 mM HEPES pH8; 100 mM KCl; 2 mM EDTA; 10% Glycerol; 0.1% NP-40; 1 mM DTT; protease inhibitors (Roche); phosphatase inhibitor cocktail 3 (MilliporeSigma)) for 30 minutes on ice. Lysates were frozen in dry ice for 5 minutes, then thawed and centrifuged for 20 minutes at 16,000xg at 4°C. An aliquot representing the input was removed before cleared supernatants were incubated with equilibrated anti-FLAG M2 Affinity Gel (MilliporeSigma) for 1-2 hours at 4°C. Beads were washed 3 times with lysis buffer before resuspension in 2x SDS-PAGE sample buffer. Samples were heated at 95°C for 5 min.

To probe PLK4 modification in Figures 7D, F and G, 350 k HEK293T cells were seeded per well of a 6-well plate and subsequently transfected with 0.67 μg Myc-PLK4 and 1 μg T7-TRIM37 construct using 3.34 μL Lipofectamine 3000/P3000. Medium was changed after 6 h and cells incubated for 48 h in total before sampling. For Figure 7G, the indicated drug was added 3 and 6 h before collection. Cells were collected directly in 2x SDS-PAGE sample buffer for Figure 7D and G. For Figure 7F, cells were collected and washed once with PBS. Cells were resuspended in a modified TNTE buffer (10 mM Tris-HCl, pH = 7.4, 100 mM NaCl, 1 mM EDTA, 1 mM DTT, 0.1% TX-100; protease inhibitors; ± phosphatase inhibitor cocktail 3) and incubated for 30 min on ice before addition of MnCl_2_ and λ-phosphatase (Bio-Rad) to the appropriate samples for 30 min at 30°C. The soluble fractions were obtained by centrifugation at 16,000 x g for 30 min. To probe for PLK4 ubiquitination, 1.5×10^6^ HEK293T cells were seeded in a 10cm dish and transfected with 2 μg Myc-PLK4, 2 μg HA-Ubiquitin and 2 μg eGFP or T7-TRIM37 with 12 μL Lipofectamine 3000/P3000. The medium was changed after 16 h and cells harvested after 48 h. Cells were washed once with PBS and resuspended in modified TNTE buffer and the soluble fraction was obtained as described above. Lysates were incubated with 3 μg anti-Myc antibodies (Table S3) and incubated for 1 h at 4°C. Equilibrated Protein G Sepharose 4 Fast Flow beads (Cytiva) were added and samples further incubated for 1 h at 4°C. Immunoprecipates were washed 3x with modified TNTE buffer and eluted by addition of 2x SDS-PAGE sample buffer and heating at 95°C for 5 min.

### FRAP analysis

For experiments using disruption lines 62.5 k cells were seeded per well of an 8-well LabTekII chamber. Cells were transfected with 400 ng pcDNA5 GFP-PLK4 or pcDNA5 GFP-PLK4 kin+L1 using 0.8 uL and 0.6 uL P3000/Lipofectamine 3000. Media was removed after 6 h and replaced with media containing DMSO or centrinone B. Cells were incubated approximately 16 h before imaging using a Nikon A1R-HD25 scanning laser confocal microscope with a LUN4 laser unit and GaAsP PMT. A single Z-slice was imaged in Galvano mode, 1.2 μs dwell time using a 488 nm excitation wavelength, a 521/42 bandpass emission filter and a 60x NA 1.2 water immersion objective. GFP-PLK4 condensates were imaged at 0, 4 and 8 s before bleaching for 500 ms using 60% 488 laser power and 16 fps scan speed. Images were acquired every 4 s for a total of 90 s after bleaching. Imaging parameters were adjusted as needed between replicates (typically 1.2% laser power with gain setting of 10). Analysis was performed using NIS-elements ‘time measurement’ module. For each image an ROI was drawn around the targeted area, a similar unbleached area and a background region. Where appropriate, the ROI was moved to track the structure of interest. The signal from the targeted area (ROI1) was background subtracted (ROI3) and then normalized using the unbleached area (ROI2) to correct for photobleaching during imaging. Ten to 15 GFP-PLK4 condensates from different cells were analyzed per condition per replicate.

## SUPPLEMENTAL FIGURE LEGENDS

**Figure 1 – figure supplement 1.**
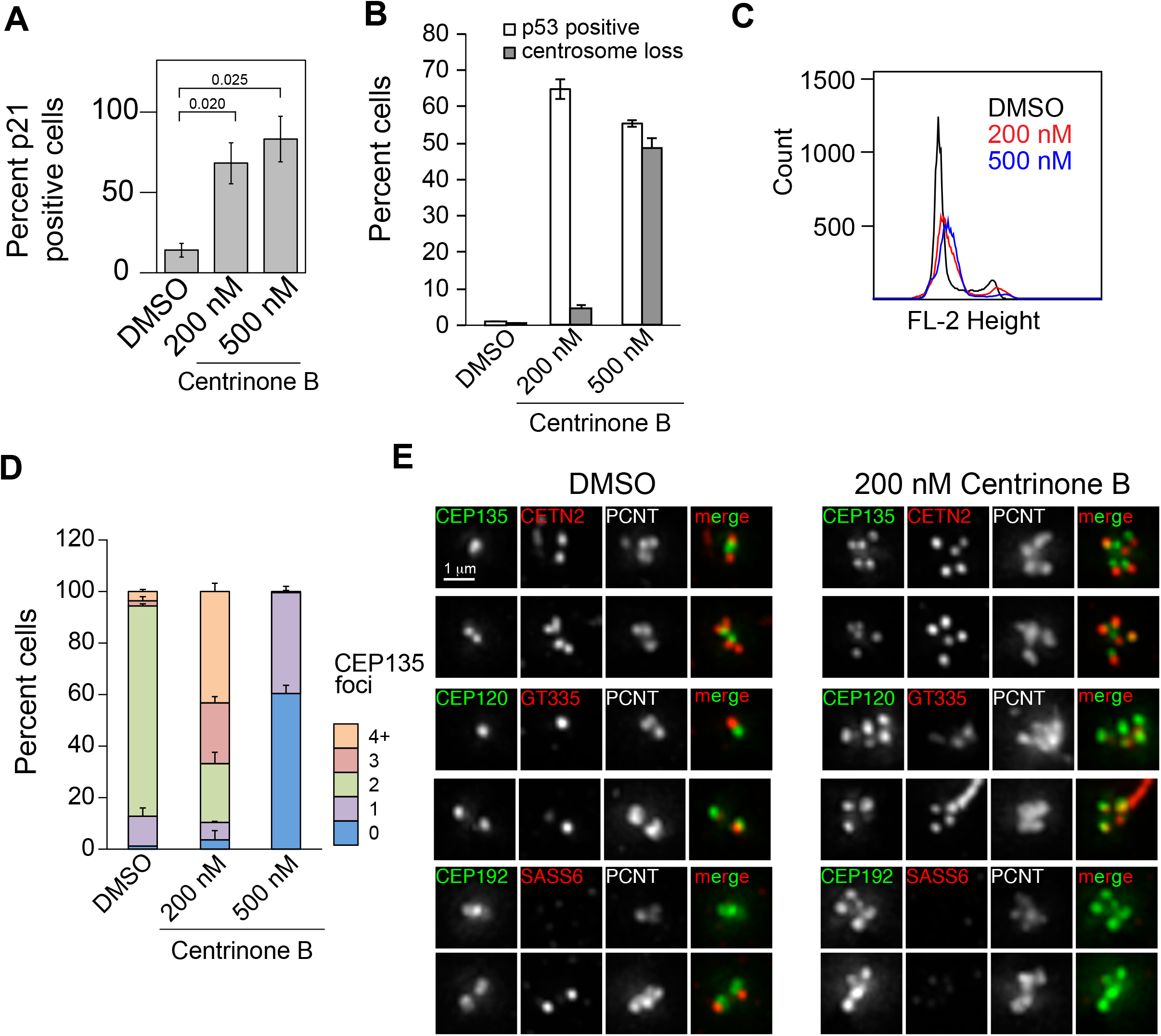
Characterization of growth arrest and centriole abnormalities after PLK4 inhibition. **(A)** RPE-1 cells were treated with DMSO, or 200 or 500 nM centrinone B for 4 days. Fixed cells were immunostained for p21 and the percent p21-positive cells was quantified. Mean and standard deviation shown (n = 2, N ≥ 100). **(B)** Cells prepared as in Figure 1C were automatically analyzed for nuclear p53 or manually scored for the presence or absence of CEP135 foci. Mean and standard deviation shown (n = 3, N ≥ 100). **(C)** RPE-1 cells treated as in (A) were fixed, stained with propidium iodide and subjected to flow cytometry to analyze DNA content. The results from a representative experiment are shown. **(D)** Cells prepared as in Figure 1D were manually scored for CEP135 foci. Means and standard deviation shown (n = 3, N ≥ 100). **(E)** RPE-1 cells treated with DMSO or 200 nM centrinone B for 4 days were fixed and stained for the indicated proteins. Two representative images per condition are shown.

**Figure 2 – figure supplement 1.**
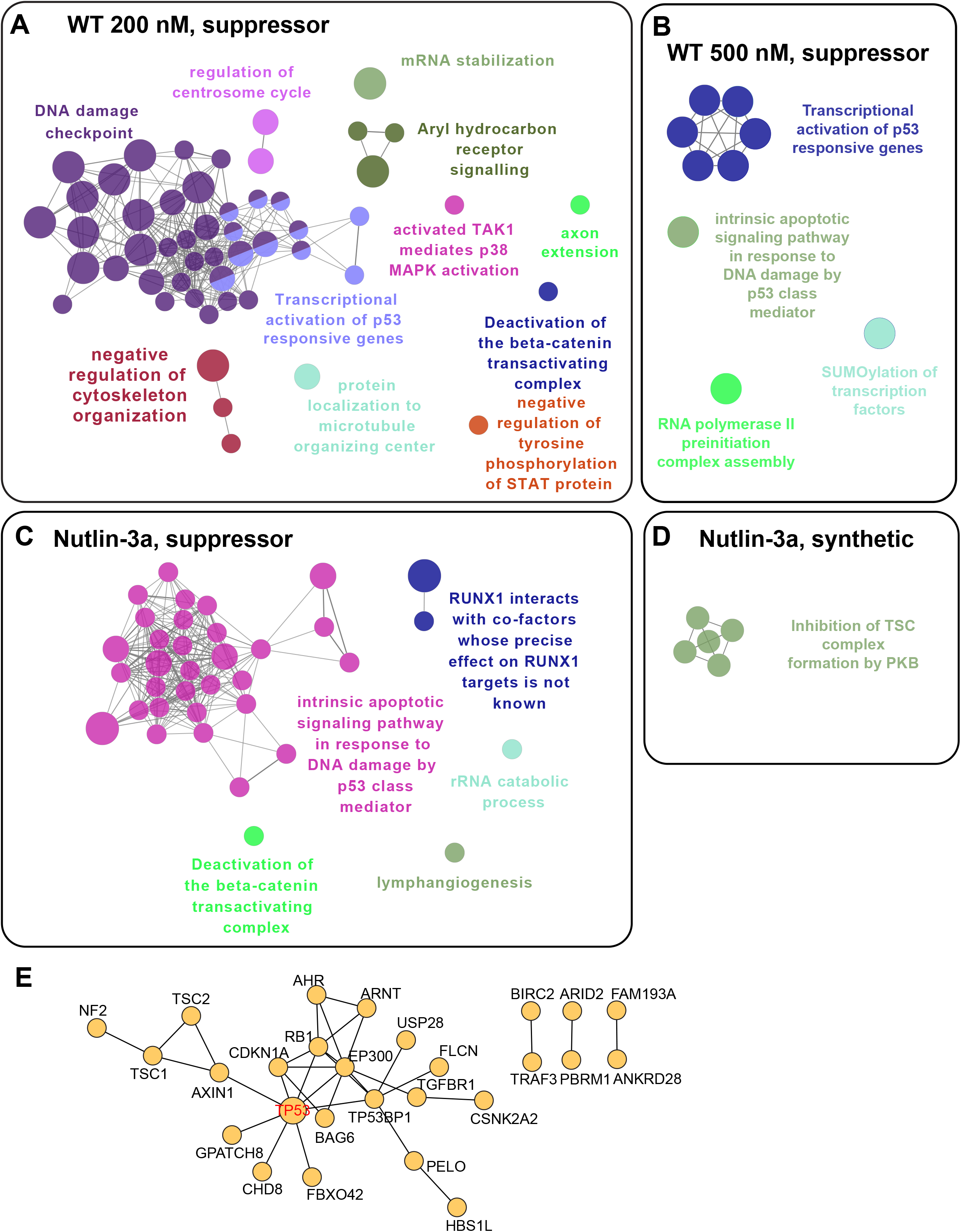
Bioinformatic analysis of CRISPR/Cas9 screens. **(A-D)** Biological process enrichment from significant genes scoring in each screen using RPE-1 cells. Screen type indicated in each figure. **(E)** Protein-protein interaction network of genes identified as suppressors in the Nutlin-3a screen.

**Figure 2 – figure supplement 2.**
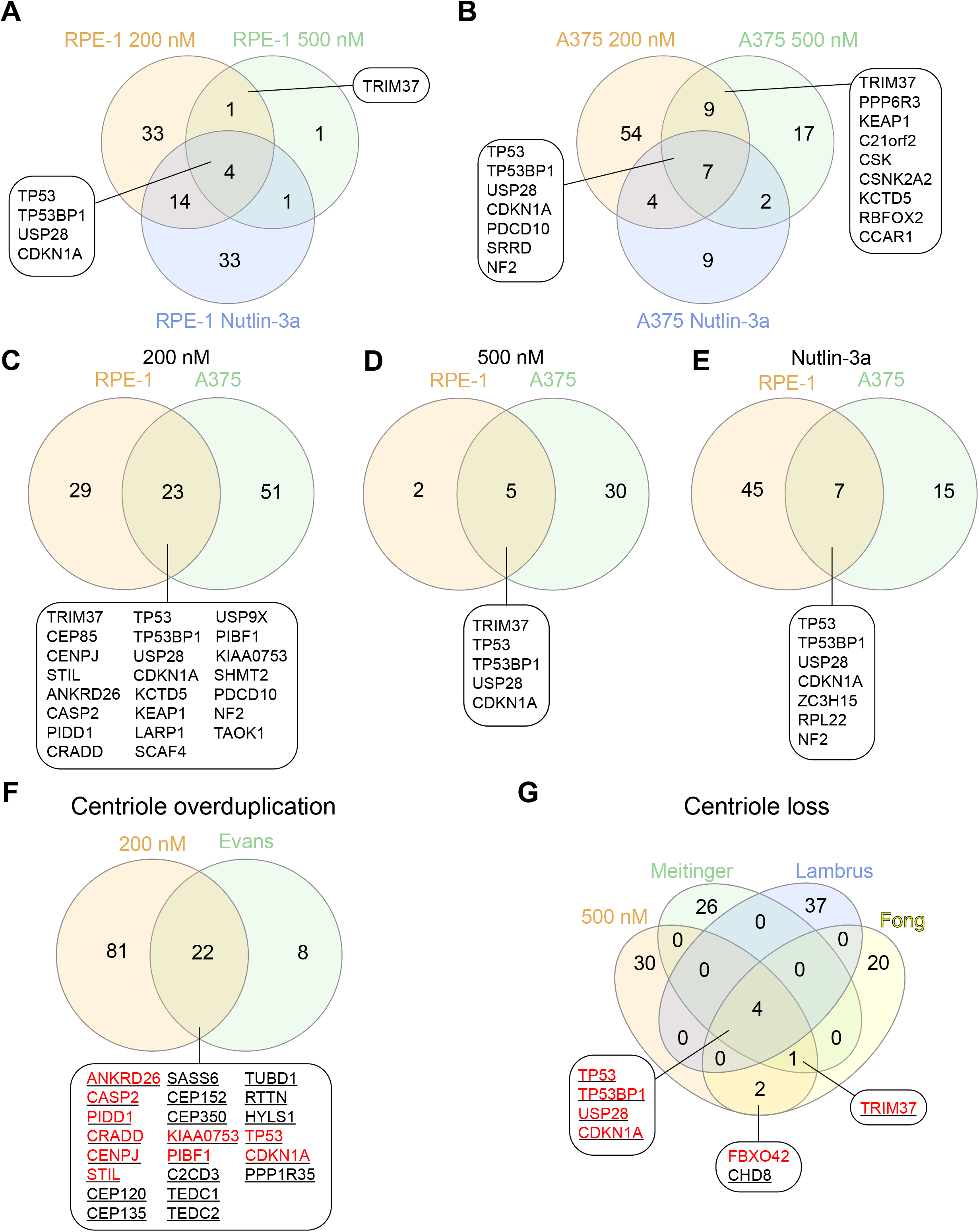
Comparison of PLK4 screens. **(A-E)** Venn diagrams showing the overlap among the hits from screens performed in this study. Screen conditions indicated in each panel. Genes from selected overlap regions are shown. **(F)** Comparison of genes identified using 200 nM centrinone B in both RPE-1 and A375 (200 nM) and after PLK4 overexpression in RPE-1 cells (Evans). Genes identified in RPE-1 cells coloured red, genes identified in A375 cells underlined. **(G)** Comparison of genes from 500 nM screens performed here in RPE-1 and A375 cells (500 nM) with other PLK4 inhibition screens performed using RPE-1 cells (Meitinger, Lambrus and Fong). Colour scheme identical to panel F.

**Figure 3 – figure supplement 1.**
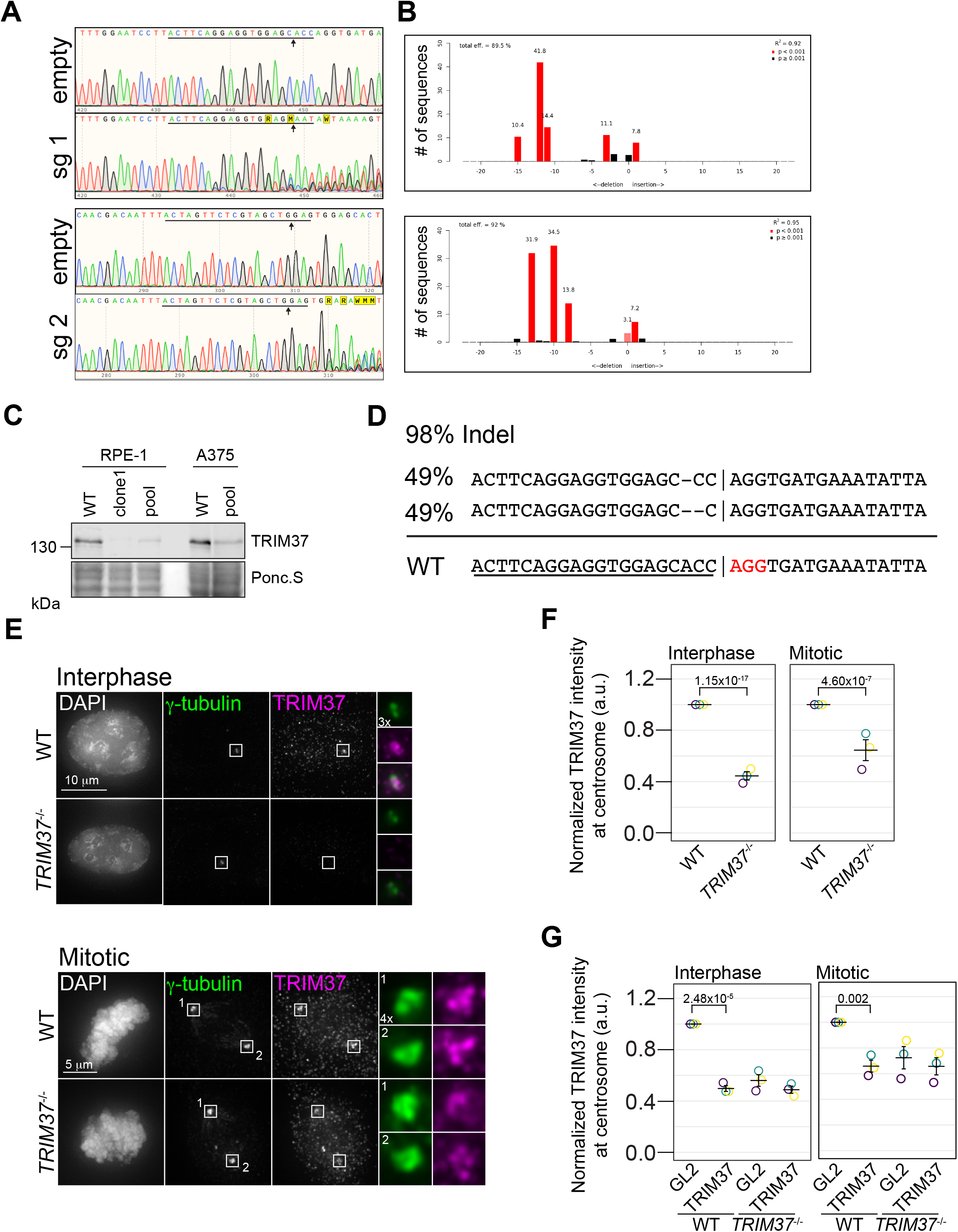
Characterization of TRIM37 gene disrupted lines. **(A)** Samples from control RPE-1 Cas9 (empty) or cells expressing the indicated TRIM37 sgRNA were collected and subjected to Sanger sequencing. Underline indicates sgRNA position and arrow indicates predicted Cas9 cleavage site. **(B)** Sequences from (A) were analyzed using CRISPR TIDE. Efficiency and indel distribution are shown. **(C)** The indicated cell lines were analyzed by Western blotting for TRIM37. ‘Clone 1’ indicates single RPE-1 *TRIM37*^−/-^ clonal line. ‘Pool’ represents RPE-1 and A375 *TRIM37*^−/-^ pools. Ponc.S indicates total protein. **(D)** Genomic DNA from clonal RPE-1 *TRIM37*^−/-^ gene disruption analyzed using Synthego CRISPR analysis tool. WT sequence is shown at bottom. sgRNA sequence is underlined and PAM coloured red. Vertical line indicates predicted Cas9 cleavage site. The deconvolved sequences indicate alleles with 1 and 2 nucleotide deletions. **(E)** RPE-1 and RPE-1 *TRIM37*^−/-^ cells were pre-extracted and fixed for immunofluorescence using the indicated antibodies. Insets enlarged 3x and 4x as indicated. **(F)** TRIM37 intensity at the centrosome was automatically determined in interphase and mitotic cells. (interphase; n = 3, N ≥ 209, mitotic; n = 3, N = 30). **(G)** RPE-1 and RPE-1 *TRIM37*^−/-^ cells were treated with control siRNA (GL2), or siRNA directed against TRIM37. The centrosomal intensity of TRIM37 was determined in interphase and mitotic cells. (interphase; n = 3, N ≥ 179, mitotic n = 3, N ≥ 30). Significant *p*-values from a pairwise test comparing control siRNA to TRIM37 siRNA are shown.

**Figure 3 – figure supplement 2.**
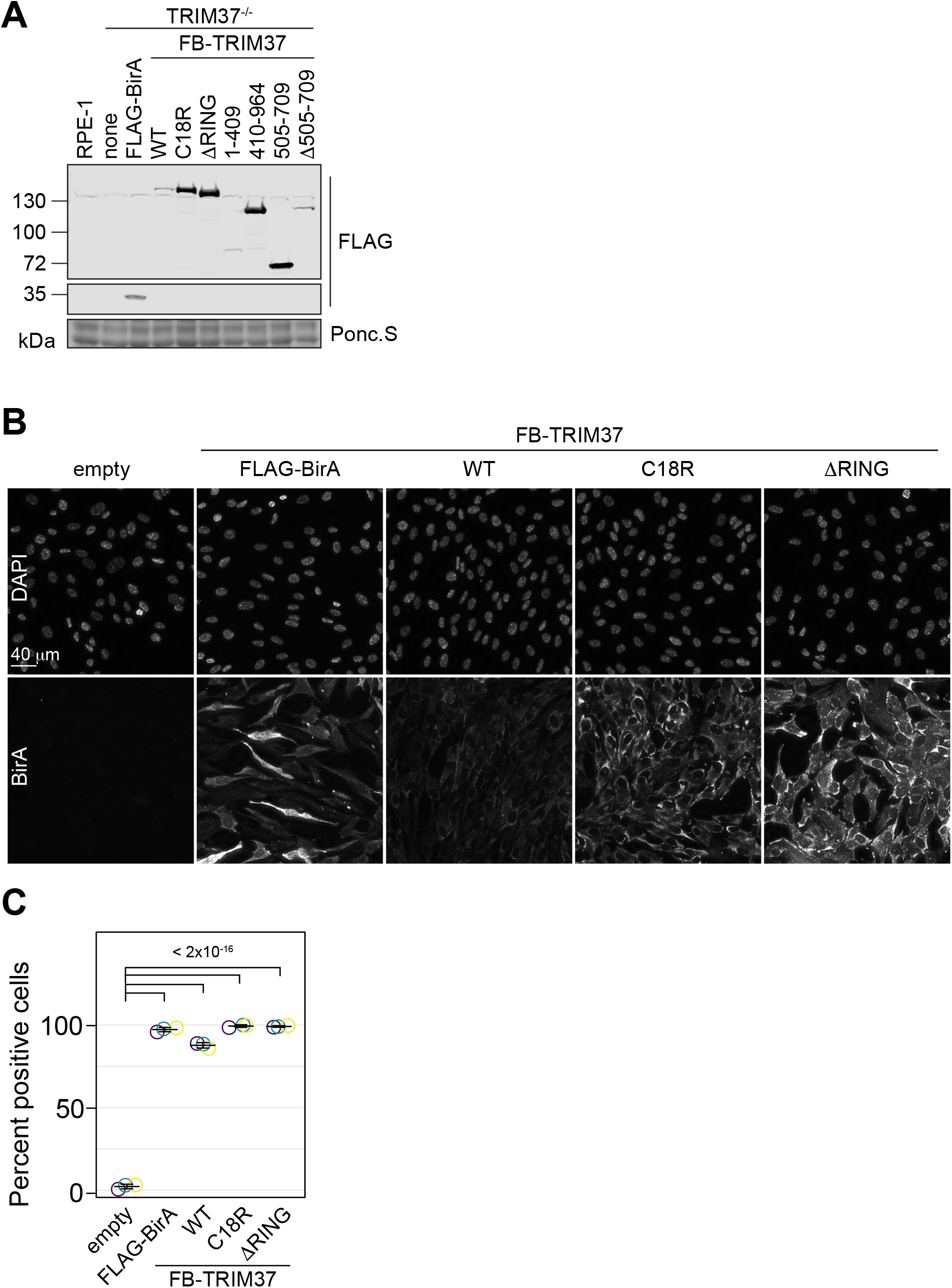
Characterization of clonal *TRIM37*^−/-^ rescue lines. **(A)** WT RPE-1 or TRIM37-/- cells stably expressing FLAG-BirA or the indicated FB-TRIM37 mutant were processed for Western blot using an anti-FLAG antibody. Ponc.S indicates total protein. **(B)** RPE-1 *TRIM37*^−/-^ cells stably expressing FLAG-BirA or the indicated FB-TRIM37 mutant were fixed and stained for BirA. **(C)** Quantification of cells in (B). The number of cells with a fluorescence intensity above a threshold were scored as positive (n = 3, N ≥ 240). Significant *p*-values from Dunnett post-hoc test using ‘empty’ as control after one-way ANOVA shown.

**Figure 4 – figure supplement 1.**
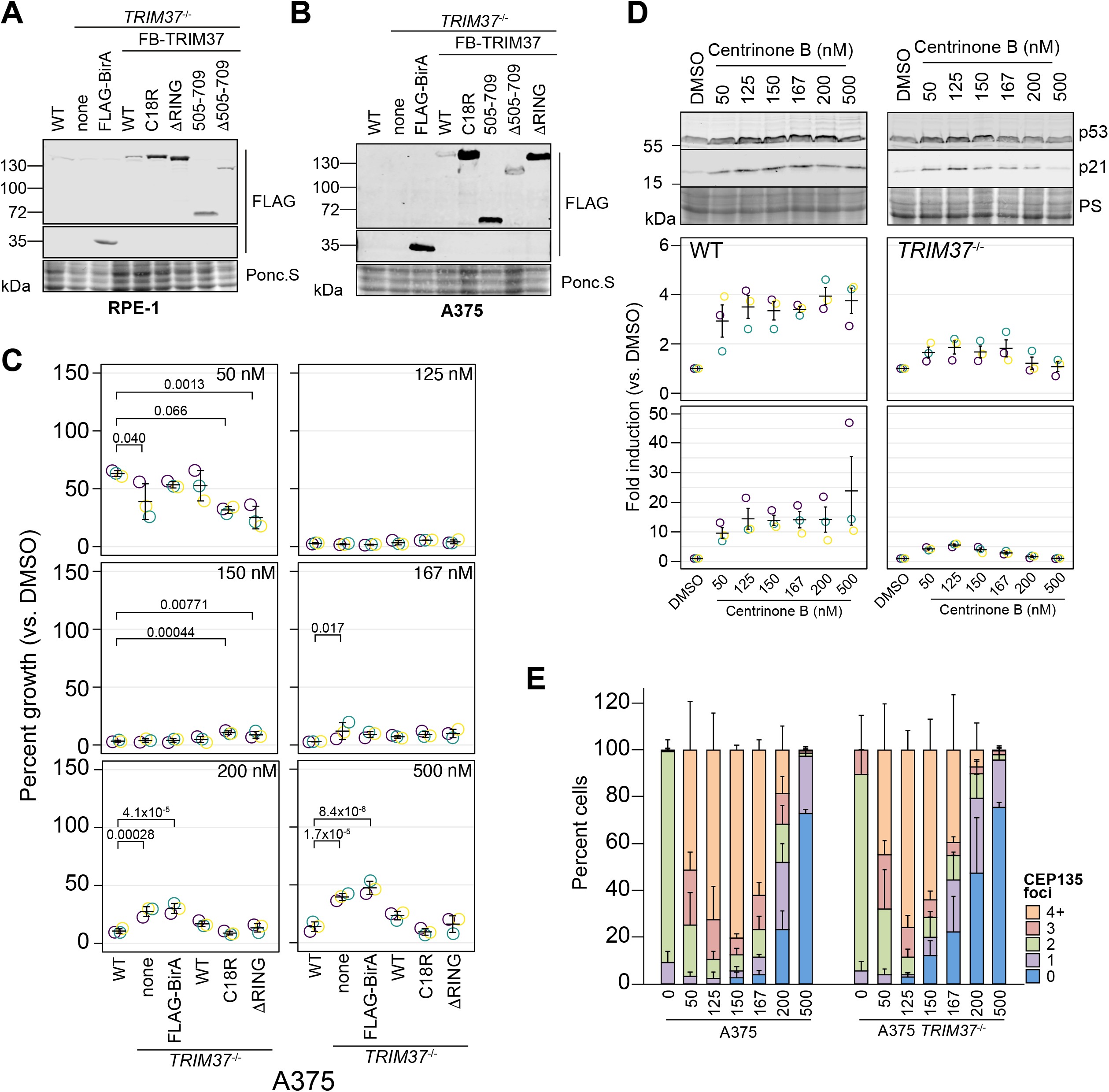
Cellular response to varying centrinone B treatments. **(A)** WT RPE-1 and a pooled RPE-1 *TRIM37*^−/-^ line stably expressing FLAG-BirA of the indicated FB-TRIM37 fusion protein were harvested and analyzed by Western blot using an anti-FLAG antibody. Ponc.S indicates total protein. **(B)** WT A375 and a pooled A375 *TRIM37*^−/-^ line stably expressing FLAG-BirA of the indicated FB-TRIM37 fusion protein were harvested and analyzed by Western blot using an anti-FLAG antibody. Ponc.S indicates total protein. **(C)** WT A375, *TRIM37*^−/-^ (pool) and *TRIM37*^−/-^ expressing FB or the indicated FB-TRIM37 construct were seeded for clonogenic assays and grown in DMSO or the indicated concentration of centrinone B for 14 days. Colony density was quantified and growth compared to that in DMSO determined. Means from each replicate are shown as open circles. Resulting mean and standard deviation shown (n = 3). Significant *p*-values from Dunnett post-hoc test using ‘WT’ as control after one-way ANOVA shown. **(D)** WT RPE-1 or *TRIM37*^−/-^ cells were treated with DMSO or the indicated concentration of centrinone B for 4 d. Samples were prepared for Western blot and probed for p53 and p21. PS indicates total protein. Band intensity was determined and normalized to DMSO for each cell line. Means from each replicate are shown as open circles. Resulting mean and standard deviation shown (n = 3). **(E)** WT or *TRIM37*^−/-^ (pool) A375 cells were treated with the indicated concentration of centrinone B (nM) for 4 d before fixing and staining for CEP135. CEP135 foci per cell were manually counted. Mean and standard deviation shown (n = 3, N ≥ 63 per condition).

**Figure 4 – figure supplement 2.**
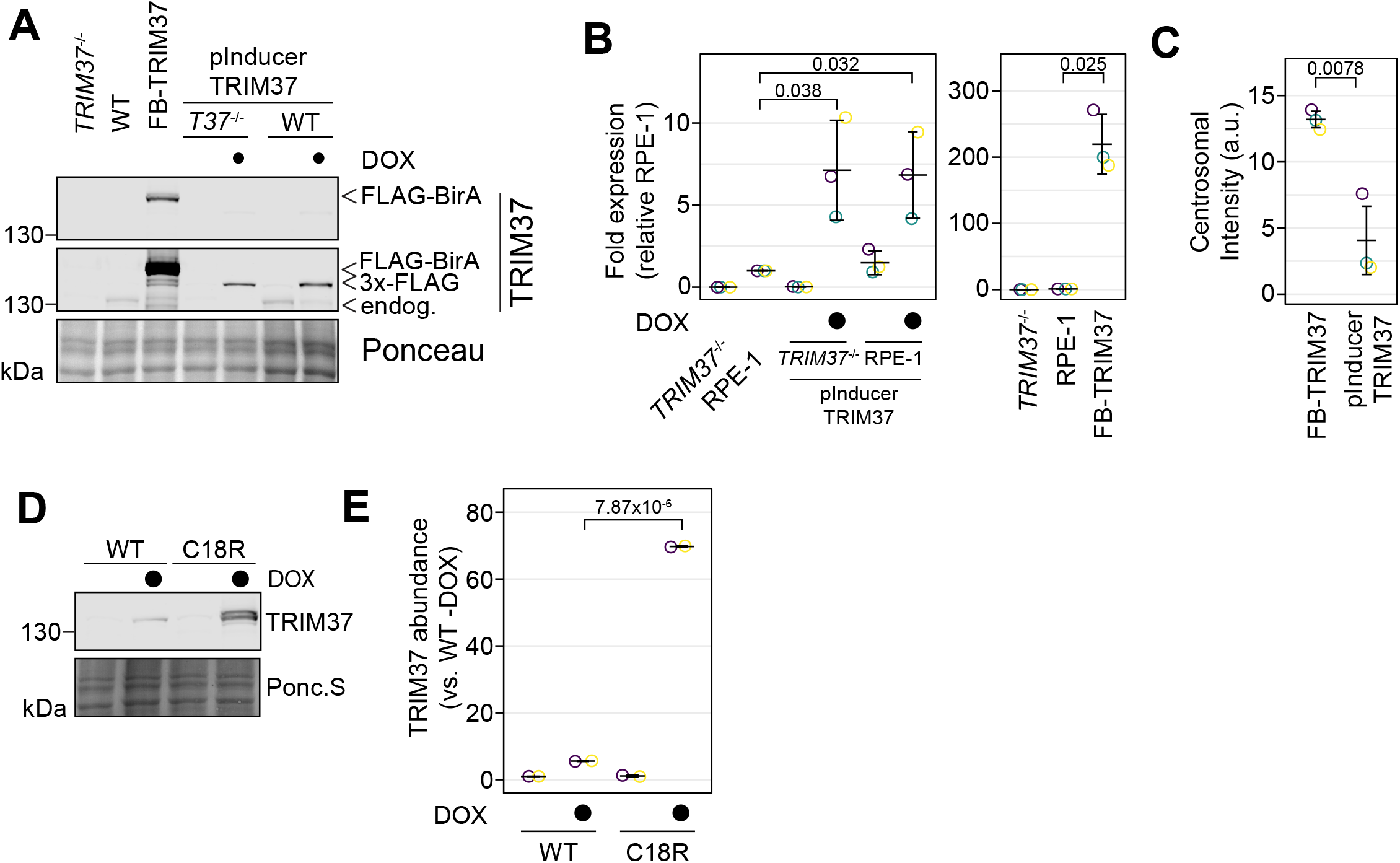
Comparison of protein abundance in stable and inducible cell lines. **(A)** The indicated cell lines were grown in the absence or presence of doxycycline for 3 d (*TRIM37*^−/-^: RPE-1 *TRIM37*^−/-^; WT:RPE-1, FB-TRIM37: FB-TRIM37 expressed in *TRIM37*^−/-^, pInducer TRIM37 expressed in WT: RPE-1 and *T37*^−/-^: RPE-1 *TRIM37*^−/-^). Cells were prepared for Western blot and probed for TRIM37. Ponc.S indicates total protein. **(B)** Quantification of (A). Relative intensities from each replicate are shown as open circles. Resulting mean and standard deviation shown (n = 3). Significant *p*-values from Dunnett post-hoc test using ‘RPE1’ as control after one-way ANOVA shown. Only RPE-1, *TRIM37*^−/-^ pInducer TRIM37 +DOX and RPE-1 pInducer TRIM37 +DOX were compared. RPE-1 and FB-TRIM37 were compared using a pairwise t-test. **(C)** The indicated cell lines from (A) were also fixed and stained for TRIM37 and γ-tubulin. The total centrosomal intensity in the TRIM37 channel was determined. Means from each replicate are shown as open circles. Resulting mean and standard deviation shown (n = 3, N ≥ 164). Significant *p*-value from a pairwise t-test is shown. **(D)** RPE-1 *TRIM37*^−/-^ cells expressing inducible WT or C18R TRIM37 were grown in the absence or presence of doxycycline for 3 d. Samples were processed for Western blot and probed for TRIM37. Ponc. S indicates total protein. **(E)** Quantification of (D). Relative intensities from each replicate are shown as open circles. Resulting mean and standard deviation shown (n = 3). The doxycycline induced samples were compared using a pairwise t-test.

**Figure 4 – figure supplement 3.**
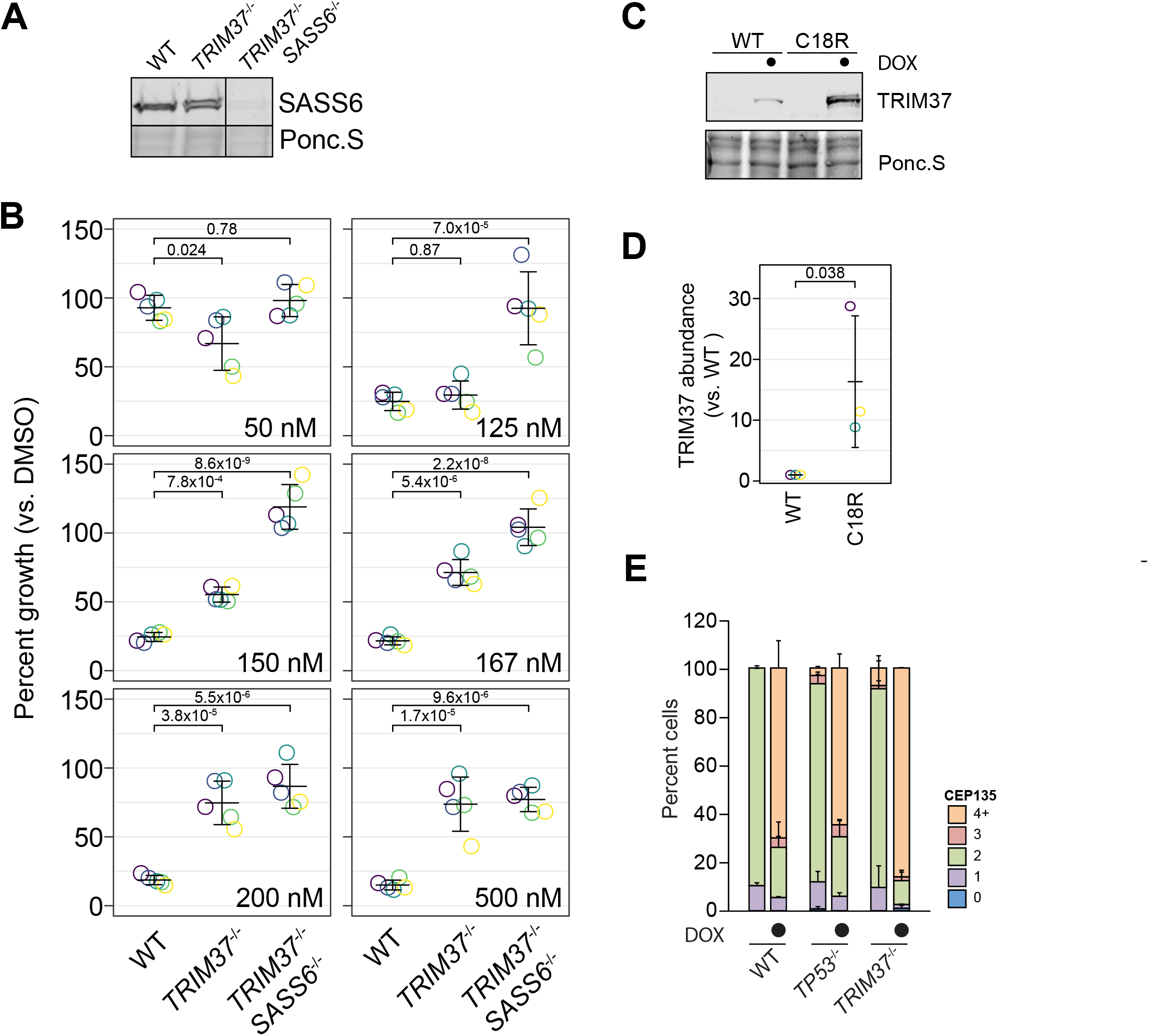
Characterization of *SASS6*^−/-^ and pInducer PLK4 cell lines. **(A)** WT RPE-1, *TRIM37*^−/-^ and *TRIM37*^−/-^ *SASS6*^−/-^ cells were prepared for Western blot and probed for SASS6. Ponc.S indicates total protein. **(B)** WT RPE-1, *TRIM37*^−/-^ and *TRIM37*^−/-^/*SASS6*^−/-^ cells were seeded for clonogenic assays and grown in DMSO or the indicated concentration of centrinone B for 14 days. Colony density was quantified and growth compared to that in DMSO determined. Means from each replicate are shown as open circles. Resulting mean and standard deviation shown (n = 5). Significant *p*-values from Dunnett post-hoc test using ‘WT’ as control after one-way ANOVA shown. **(C)** RPE-1 *TRIM37*^−/-^/*SASS6*^−/-^ cells expressing inducible WT or C18R TRIM37 were grown in the absence or presence of doxycycline for 3 d. Samples were prepared for Western blot and probed for TRIM37. Ponc.S indicates total protein. **(D)** Quantification of the induced samples in (C). WT and C18R were compared using a pairwise t-test. **(E)** WT RPE-1, *TP53*^−/-^ and *TRIM37*^−/-^ expressing inducible PLK4-3xFLAG were grown in the absence and presence of doxycycline for 16 h. Cells were fixed and stained for CEP135. The number of CEP135 foci was manually counted. Means and standard deviation are shown (n = 2, N ≥ 54 per condition).

**Figure 6 – figure supplement 1.**
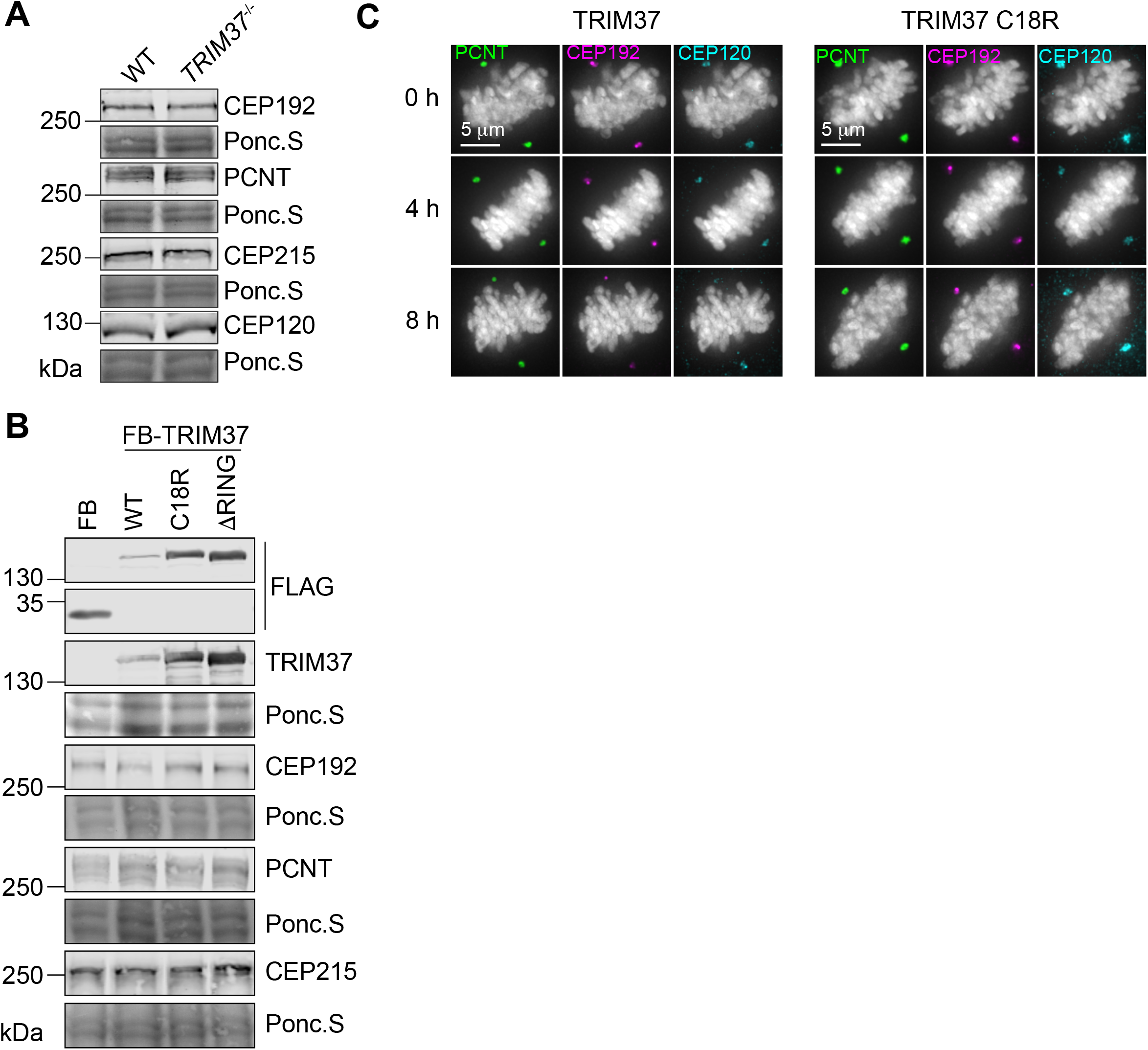
TRIM37 negatively regulates PCM and centriole proteins. **(A)** WT RPE-1 and *TRIM37*^−/-^ cells were processed for Western blot and probed for the indicated protein. Ponc.S indicates total protein. Quantification in Figure 6A. **(B)** *TRIM37*^−/-^ RPE-1 cells stably expressing FLAG-BirA (FB) or the indicated FB-TRIM37 protein were processed for Western blot and probed for the indicated protein. Ponc.S indicates total protein. Quantification in Figure 6B. **(C)** RPE-1 *TRIM37*^−/-^ cells expressing inducible TRIM37 or TRIM37 C18R were induced with doxycycline for the indicated time. Cells were fixed and stained for the indicated protein. Sample images of mitotic cells quantified in Figure 6D.

**Figure 6 – figure supplement 2.**
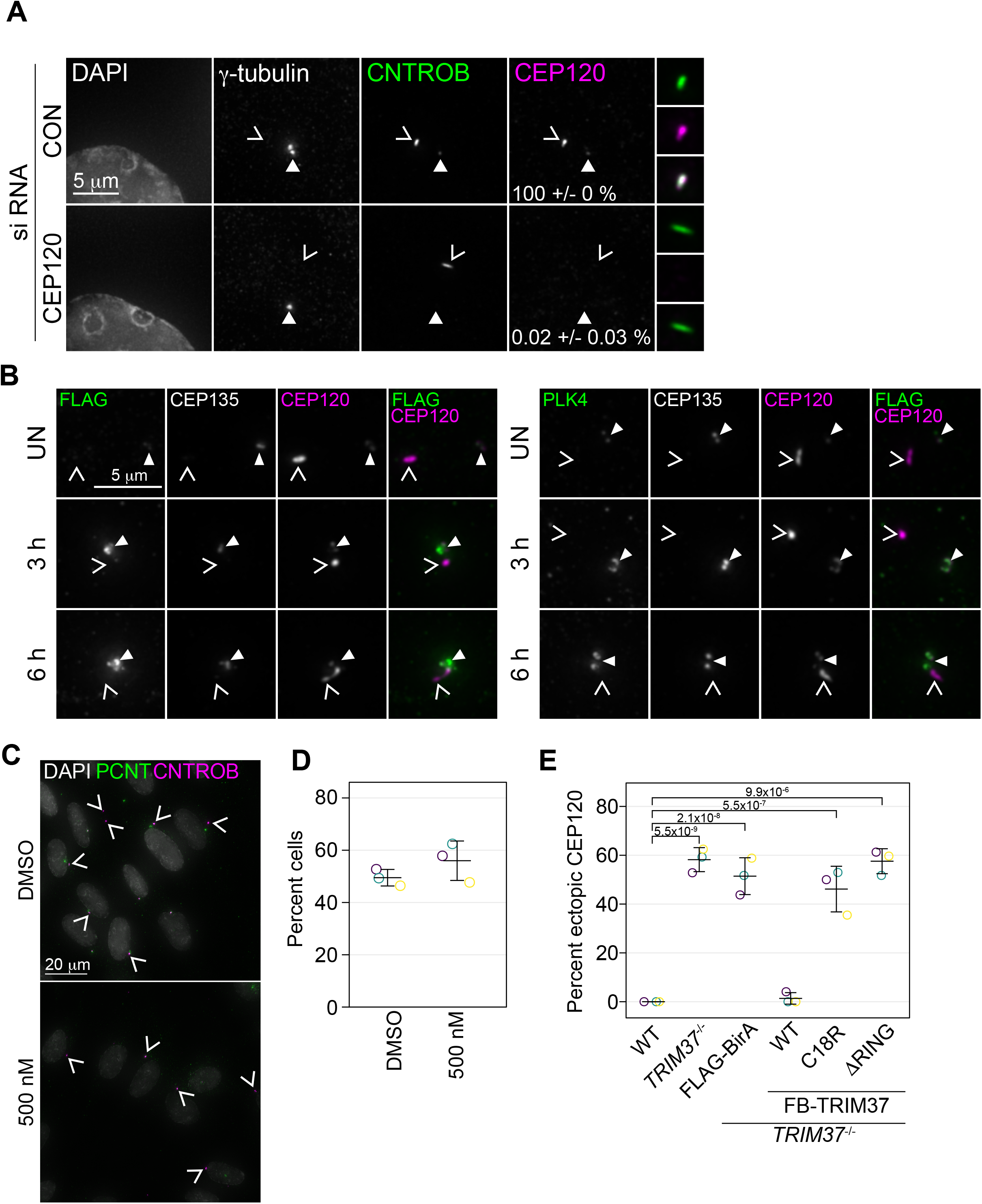
Characterization of ectopic centrosomal aggregates. **(A)** RPE-1 cells were treated with non-targeting (CON) siRNA or siRNA targeting CEP120 (CEP120). Cells were fixed after 72 h and stained for the indicated protein. Closed arrowhead indicates centrosome based on γ-tubulin. Caret mark (open arrowhead) indicates ectopic structures based on CEP120 and/or CNTROB. Cells were manually scored for the number of cells with ectopic CNTROB structures that also co-stained with CEP120 (mean with standard deviation). **(B)** *TRIM37*^−/-^ RPE-1 cells expressing inducible PLK4-3xFLAG were treated with doxycycline. Cells were harvested at the indicated time, fixed and stained for FLAG (left panels) or PLK4 (right panels) and the indicated proteins. Closed arrowhead indicates the centrosome based on CEP135 and the caret mark indicates ectopic structures based on the non-centrosomal CEP120 signal. **(C)** *TRIM37*^−/-^ RPE-1 cells were grown in the absence and presence of 500 nM centrinone B for 3 d. Cells were fixed and stained for the indicated proteins. **(D)** Quantification of cells in (C). The number of cells with non-centrosomal CNTROB was scored. Means and standard deviation shown. Control samples were compared to centrinone B-treated using a pairwise t-test and no significant difference was observed.

**Figure 7 – figure supplement 1.**
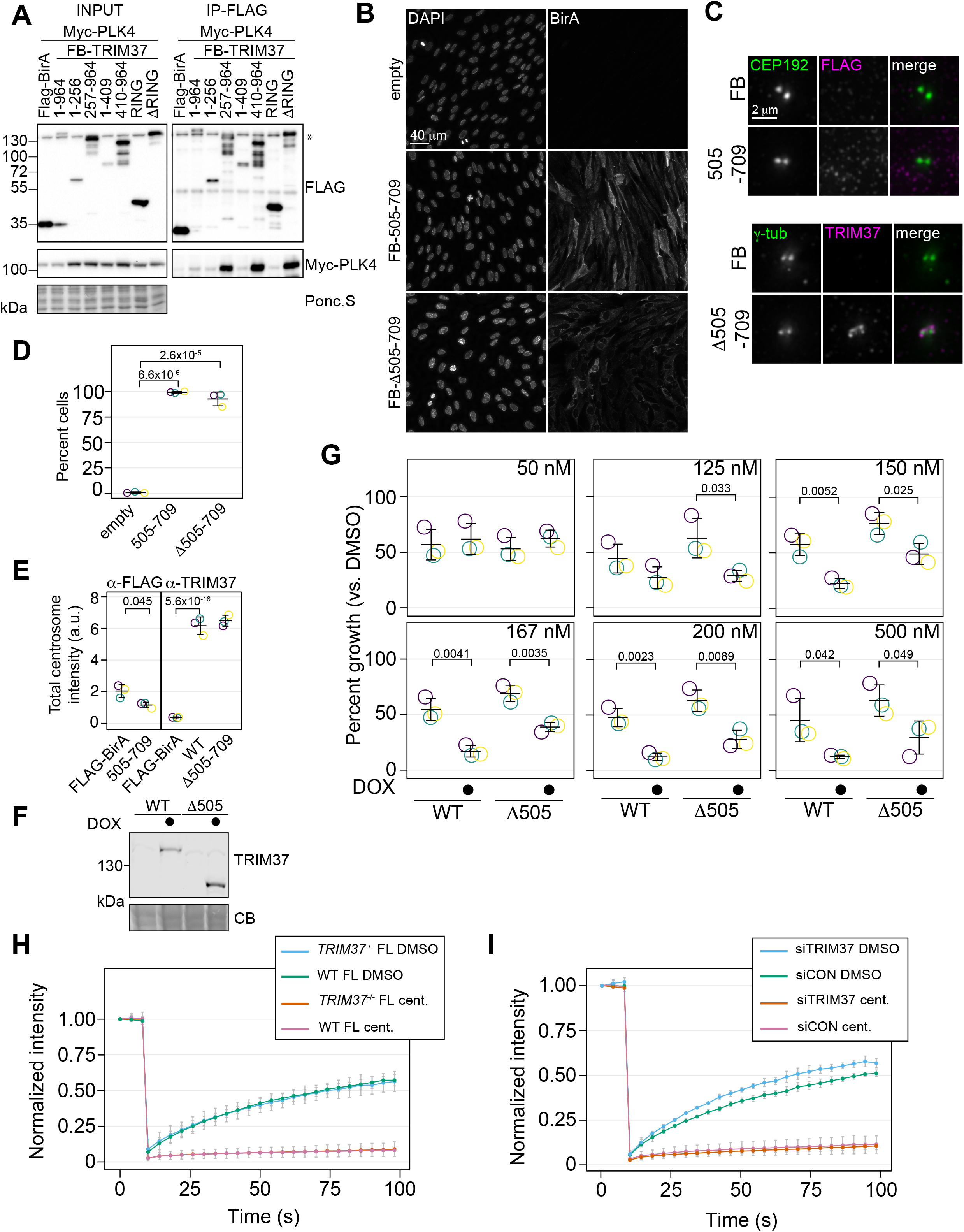
Characterization of TRIM37 interactions with PLK4. **(A)** RPE-1cells were transiently transfected to express Myc-PLK4 and FLAG-BirA or the indicated FB-TRIM37 construct (top). Cells were lysed and subjected to anti-FLAG immunoprecipitation. Input and immunoprecipitates were analyzed by immunoblotting with the indicated antibodies. Ponc.S indicates total protein. * indicates position of FLAG-Cas9. **(B)** RPE-1 *TRIM37*^−/-^ cells stably expressing the indicated FB-TRIM37 protein were fixed and stained using an anti-BirA antibody. **(C)** Cells as in (B) were pre-extracted, fixed and stained for the indicated protein (FB: FLAG-BirA). **(D)** Quantification of cells in (B). Means from each replicate are shown as open circles. Resulting mean and standard deviation shown (n = 3, N ≥ 296). Significant *p*-values from Dunnett post-hoc test using ‘empty’ as a control after a one-way ANOVA are shown. **(E)** Quantification of cells in (C). Means from each replicate are shown as open circles. Resulting mean and standard deviation shown (n = 3, N ≥ 78). Samples stained with α-FLAG were compared using a pairwise t-test. For samples stained with α-TRIM37, significant *p*-values from Dunnett post-hoc test using ‘WT’ as a control after a one-way ANOVA are shown. Note that the results from the α-TRIM37 samples and those in Figure 3I and J are from the same experiment therefore ‘FLAG-BirA’ and ‘WT’ are duplicated in these panels. **(F)** RPE-1 *TRIM37*^−/-^ cells expressing the inducible FB-TRIM37 protein indicated were grown for 3 d in the absence and presence of doxycycline. Samples were prepared for Western blot and probed for TRIM37. CB indicates total protein. **(G)** RPE-1 *TRIM37*^−/-^ cells expressing the indicated inducible TRIM37 construct were seeded for clonogenic assays in the absence and presence of doxycycline and DMSO or the indicated concentration of centrinone B. After incubation for 14 d, colony density was quantified and growth compared to that in DMSO determined. Means from each replicate are shown as open circles. Resulting mean and standard deviation shown (n = 3). Significant *p*-values from pairwise t-tests comparing −DOX and +DOX for each line are shown. **(H)** WT RPE-1 and *TRIM37*^−/-^ cells were transfected to express GFP-PLK4. FRAP analyses were performed after treating cells with DMSO or 500 nM centrinone B (cent.) for 16 h. The mean and standard deviation among the independent replicates is shown (n = 3, N ≥ 10). **(I)** WT RPE-1 cells were treated with control siRNA (siCON) or siRNA against TRIM37 (siTRIM37) for 72 h. Cells were transfected to express GFP-PLK4 after 48 h and further treated with DMSO or 500 nM centrinone B (cent.) for the final 16 h before performing FRAP analyses. The mean and standard deviation among the independent replicates is shown (n = 3, N ≥ 10).

**Figure 7 – figure supplement 2.**
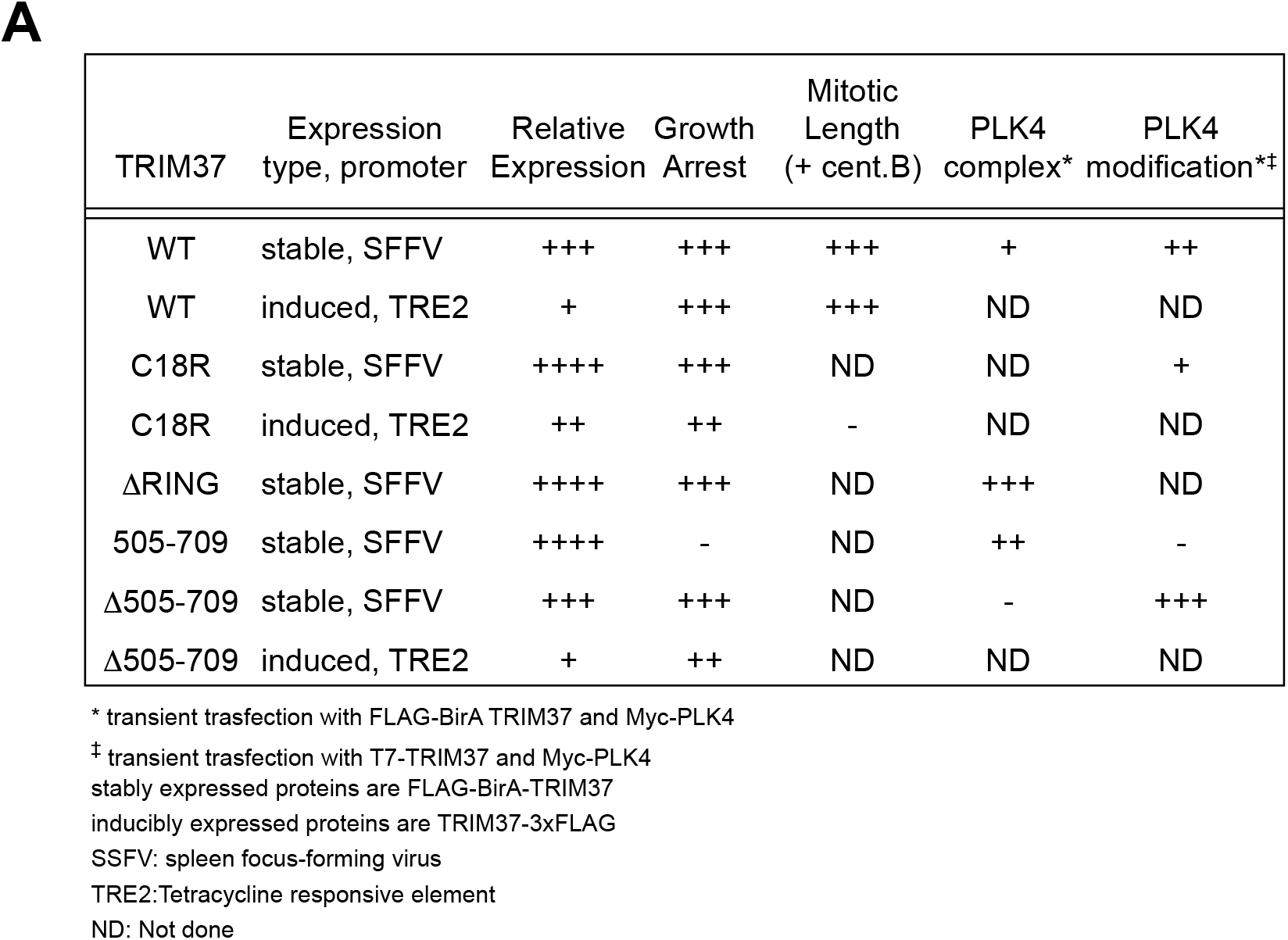
Summary of TRIM37 rescue constructs used and resulting phenotypes. **(A)** Table summarizing TRIM37 rescue constructs, expression system and relative phenotypes.

## SUPPLEMENTAL FILES

**Table S1. Summary of screening data. (Excel file)**

**Table S2. Gene enrichment details for RPE-1 screens. (Excel file)**

**Table S3. Reagents used in this study. (Excel file)**

**Source_data. All raw data used for quantification. (Excel file)**

**Western_raw. All unaltered original images for Western blot. (PDF file)**

## REFERENCES

1. Conduit, P.T., A. Wainman, and J.W. Raff, Centrosome function and assembly in animal cells. Nat Rev Mol Cell Biol, 2015. 16(10): p. 611–24.

2. Gonczy, P. and G.N. Hatzopoulos, Centriole assembly at a glance. J Cell Sci, 2019. 132(4).

3. Fode, C., C. Binkert, and J.W. Dennis, Constitutive expression of murine Sak-a suppresses cell growth and induces multinucleation. Mol Cell Biol, 1996. 16(9): p. 4665–72.

4. Sonnen, K.F., et al., Human Cep192 and Cep152 cooperate in Plk4 recruitment and centriole duplication. J Cell Sci, 2013. 126(Pt 14): p. 3223–33.

5. Kim, T.S., et al., Hierarchical recruitment of Plk4 and regulation of centriole biogenesis by two centrosomal scaffolds, Cep192 and Cep152. Proc Natl Acad Sci U S A, 2013. 110(50): p. E4849–57.

6. Moyer, T.C., et al., Binding of STIL to Plk4 activates kinase activity to promote centriole assembly. J Cell Biol, 2015. 209(6): p. 863–78.

7. Ohta, M., et al., Direct interaction of Plk4 with STIL ensures formation of a single procentriole per parental centriole. Nat Commun, 2014. 5: p. 5267.

8. Liu, Y., et al., Direct binding of CEP85 to STIL ensures robust PLK4 activation and efficient centriole assembly. Nat Commun, 2018. 9(1): p. 1731.

9. Schmidt, T.I., et al., Control of centriole length by CPAP and CP110. Curr Biol, 2009. 19(12): p. 1005–11.

10. Kohlmaier, G., et al., Overly long centrioles and defective cell division upon excess of the SAS-4-related protein CPAP. Curr Biol, 2009. 19(12): p. 1012–8.

11. Tang, C.J., et al., CPAP is a cell-cycle regulated protein that controls centriole length. Nat Cell Biol, 2009. 11(7): p. 825–31.

12. Comartin, D., et al., CEP120 and SPICE1 cooperate with CPAP in centriole elongation. Curr Biol, 2013. 23(14): p. 1360–6.

13. Azimzadeh, J., et al., hPOC5 is a centrin-binding protein required for assembly of full-length centrioles. J Cell Biol, 2009. 185(1): p. 101–14.

14. Rogers, G.C., et al., The SCF Slimb ubiquitin ligase regulates Plk4/Sak levels to block centriole reduplication. J Cell Biol, 2009. 184(2): p. 225–39.

15. Cunha-Ferreira, I., et al., The SCF/Slimb ubiquitin ligase limits centrosome amplification through degradation of SAK/PLK4. Curr Biol, 2009. 19(1): p. 43–9.

16. Guderian, G., et al., Plk4 trans-autophosphorylation regulates centriole number by controlling betaTrCP-mediated degradation. J Cell Sci, 2010. 123(Pt 13): p. 2163–9.

17. Chan, J.Y., A clinical overview of centrosome amplification in human cancers. Int J Biol Sci, 2011. 7(8): p. 1122–44.

18. Ganem, N.J., S.A. Godinho, and D. Pellman, A mechanism linking extra centrosomes to chromosomal instability. Nature, 2009. 460(7252): p. 278–82.

19. Silkworth, W.T., et al., Multipolar spindle pole coalescence is a major source of kinetochore mis-attachment and chromosome mis-segregation in cancer cells. PLoS One, 2009. 4(8): p. e6564.

20. Arnandis, T., et al., Oxidative Stress in Cells with Extra Centrosomes Drives Non-Cell-Autonomous Invasion. Dev Cell, 2018. 47(4): p. 409–424 e9.

21. Holland, A.J., et al., The autoregulated instability of Polo-like kinase 4 limits centrosome duplication to once per cell cycle. Genes Dev, 2012. 26(24): p. 2684–9.

22. Lambrus, B.G., et al., A USP28-53BP1-p53-p21 signaling axis arrests growth after centrosome loss or prolonged mitosis. J Cell Biol, 2016. 214(2): p. 143–53.

23. Wong, Y.L., et al., Cell biology. Reversible centriole depletion with an inhibitor of Polo-like kinase 4. Science, 2015. 348(6239): p. 1155–60.

24. Fong, C.S., et al., 53BP1 and USP28 mediate p53-dependent cell cycle arrest in response to centrosome loss and prolonged mitosis. Elife, 2016. 5.

25. Marthiens, V., et al., Centrosome amplification causes microcephaly. Nat Cell Biol, 2013. 15(7): p. 731–40.

26. Coelho, P.A., et al., Over-expression of Plk4 induces centrosome amplification, loss of primary cilia and associated tissue hyperplasia in the mouse. Open Biol, 2015. 5(12): p. 150209.

27. Vitre, B., et al., Chronic centrosome amplification without tumorigenesis. Proc Natl Acad Sci U S A, 2015. 112(46): p. E6321–30.

28. Sercin, O., et al., Transient PLK4 overexpression accelerates tumorigenesis in p53-deficient epidermis. Nat Cell Biol, 2016. 18(1): p. 100–10.

29. Hudson, J.W., et al., Late mitotic failure in mice lacking Sak, a polo-like kinase. Curr Biol, 2001. 11(6): p. 441–6.

30. Meitinger, F., et al., 53BP1 and USP28 mediate p53 activation and G1 arrest after centrosome loss or extended mitotic duration. J Cell Biol, 2016. 214(2): p. 155–66.

31. Evans, L.T., et al., ANKRD26 recruits PIDD1 to centriolar distal appendages to activate the PIDDosome following centrosome amplification. Embo j, 2021. 40(4): p. e105106.

32. Burigotto, M., et al., Centriolar distal appendages activate the centrosome-PIDDosome-p53 signalling axis via ANKRD26. Embo j, 2021. 40(4): p. e104844.

33. Bhatnagar, S., et al., TRIM37 is a new histone H2A ubiquitin ligase and breast cancer oncoprotein. Nature, 2014. 516(7529): p. 116–20.

34. Wang, W., et al., TRIM37, a novel E3 ligase for PEX5-mediated peroxisomal matrix protein import. J Cell Biol, 2017. 216(9): p. 2843–2858.

35. Zhu, H., et al., Knockdown of TRIM37 Promotes Apoptosis and Suppresses Tumor Growth in Gastric Cancer by Inactivation of the ERK1/2 Pathway. Onco Targets Ther, 2020. 13: p. 5479–5491.

36. Fu, T., et al., ASB16-AS1 up-regulated and phosphorylated TRIM37 to activate NF-kappaB pathway and promote proliferation, stemness, and cisplatin resistance of gastric cancer. Gastric Cancer, 2021. 24(1): p. 45–59.

37. Chen, S., et al., TRIM37 Mediates Chemoresistance and Maintenance of Stemness in Pancreatic Cancer Cells via Ubiquitination of PTEN and Activation of the AKT-GSK-3beta-beta-Catenin Signaling Pathway. Front Oncol, 2020. 10: p. 554787.

38. Wang, W., et al., TRIM37 deficiency induces autophagy through deregulating the MTORC1-TFEB axis. Autophagy, 2018. 14(9): p. 1574–1585.

39. Balestra, F.R., et al., Discovering regulators of centriole biogenesis through siRNA-based functional genomics in human cells. Dev Cell, 2013. 25(6): p. 555–71.

40. Balestra, F.R., et al., TRIM37 prevents formation of centriolar protein assemblies by regulating Centrobin. Elife, 2021. 10.

41. Meitinger, F., et al., TRIM37 prevents formation of condensate-organized ectopic spindle poles to ensure mitotic fidelity. J Cell Biol, 2021. 220(7).

42. Yeow, Z.Y., et al., Targeting TRIM37-driven centrosome dysfunction in 17q23-amplified breast cancer. Nature, 2020. 585(7825): p. 447–452.

43. Meitinger, F., et al., TRIM37 controls cancer-specific vulnerability to PLK4 inhibition. Nature, 2020. 585(7825): p. 440–446.

44. Kallioniemi, A., et al., Detection and mapping of amplified DNA sequences in breast cancer by comparative genomic hybridization. Proc Natl Acad Sci U S A, 1994. 91(6): p. 2156–60.

45. Mason, J.M., et al., Functional characterization of CFI-400945, a Polo-like kinase 4 inhibitor, as a potential anticancer agent. Cancer Cell, 2014. 26(2): p. 163–76.

46. Suri, A., et al., Evaluation of Protein Kinase Inhibitors with PLK4 Cross-Over Potential in a Pre-Clinical Model of Cancer. Int J Mol Sci, 2019. 20(9).

47. Hart, T., et al., High-Resolution CRISPR Screens Reveal Fitness Genes and Genotype-Specific Cancer Liabilities. Cell, 2015. 163(6): p. 1515–26.

48. Vassilev, L.T., et al., In vivo activation of the p53 pathway by small-molecule antagonists of MDM2. Science, 2004. 303(5659): p. 844–8.

49. Li, W., et al., MAGeCK enables robust identification of essential genes from genome-scale CRISPR/Cas9 knockout screens. Genome Biol, 2014. 15(12): p. 554.

50. Cuella-Martin, R., et al., 53BP1 Integrates DNA Repair and p53-Dependent Cell Fate Decisions via Distinct Mechanisms. Mol Cell, 2016. 64(1): p. 51–64.

51. Sebti, S., et al., BAT3 modulates p300-dependent acetylation of p53 and autophagy-related protein 7 (ATG7) during autophagy. Proc Natl Acad Sci U S A, 2014. 111(11): p. 4115–20.

52. Liu, L., et al., p53 sites acetylated in vitro by PCAF and p300 are acetylated in vivo in response to DNA damage. Mol Cell Biol, 1999. 19(2): p. 1202–9.

53. Li, Z., et al., The OncoPPi network of cancer-focused protein-protein interactions to inform biological insights and therapeutic strategies. Nat Commun, 2017. 8: p. 14356.

54. Tohkin, M., et al., Aryl hydrocarbon receptor is required for p300-mediated induction of DNA synthesis by adenovirus E1A. Mol Pharmacol, 2000. 58(4): p. 845–51.

55. Tong, Q., et al., MnTE-2-PyP reduces prostate cancer growth and metastasis by suppressing p300 activity and p300/HIF-1/CREB binding to the promoter region of the PAI-1 gene. Free Radic Biol Med, 2016. 94: p. 185–94.

56. Armstrong, L.C., et al., Heterozygous loss of TSC2 alters p53 signaling and human stem cell reprogramming. Hum Mol Genet, 2017. 26(23): p. 4629–4641.

57. Lee, C.H., et al., Constitutive mTOR activation in TSC mutants sensitizes cells to energy starvation and genomic damage via p53. Embo j, 2007. 26(23): p. 4812–23.

58. Franz, M., et al., GeneMANIA update 2018. Nucleic Acids Res, 2018. 46(W1): p. W60–W64.

59. Sun, L., et al., JFK, a Kelch domain-containing F-box protein, links the SCF complex to p53 regulation. Proc Natl Acad Sci U S A, 2009. 106(25): p. 10195–200.

60. Nishiyama, M., et al., CHD8 suppresses p53-mediated apoptosis through histone H1 recruitment during early embryogenesis. Nat Cell Biol, 2009. 11(2): p. 172–82.

61. Lü, Y., et al., Genome-wide CRISPR screens identify novel regulators of wild-type and mutant p53 stability. bioRxiv, 2022.

62. Fava, L.L., et al., The PIDDosome activates p53 in response to supernumerary centrosomes. Genes Dev, 2017. 31(1): p. 34–45.

63. Gupta, G.D., et al., A Dynamic Protein Interaction Landscape of the Human Centrosome-Cilium Interface. Cell, 2015. 163(6): p. 1484–99.

64. Hatakeyama, S., TRIM Family Proteins: Roles in Autophagy, Immunity, and Carcinogenesis. Trends Biochem Sci, 2017. 42(4): p. 297–311.

65. Yamamoto, S. and D. Kitagawa, Self-organization of Plk4 regulates symmetry breaking in centriole duplication. Nat Commun, 2019. 10(1): p. 1810.

66. Leidel, S., et al., SAS-6 defines a protein family required for centrosome duplication in C. elegans and in human cells. Nat Cell Biol, 2005. 7(2): p. 115–25.

67. Dammermann, A., et al., Centriole assembly requires both centriolar and pericentriolar material proteins. Dev Cell, 2004. 7(6): p. 815–29.

68. Soucy, T.A., et al., An inhibitor of NEDD8-activating enzyme as a new approach to treat cancer. Nature, 2009. 458(7239): p. 732–6.

69. Lei, Q., et al., YLT-11, a novel PLK4 inhibitor, inhibits human breast cancer growth via inducing maladjusted centriole duplication and mitotic defect. Cell Death Dis, 2018. 9(11): p. 1066.

70. Holland, A.J. and D.W. Cleveland, Polo-like kinase 4 inhibition: a strategy for cancer therapy? Cancer Cell, 2014. 26(2): p. 151–3.

71. Kodani, A., et al., SFI1 promotes centriole duplication by recruiting USP9X to stabilize the microcephaly protein STIL. J Cell Biol, 2019. 218(7): p. 2185–2197.

72. Oliver, T.G., et al., Caspase-2-mediated cleavage of Mdm2 creates a p53-induced positive feedback loop. Mol Cell, 2011. 43(1): p. 57–71.

73. Lambrus, B.G., et al., p53 protects against genome instability following centriole duplication failure. J Cell Biol, 2015. 210(1): p. 63–77.

74. Basto, R., et al., Centrosome amplification can initiate tumorigenesis in flies. Cell, 2008. 133(6): p. 1032–42.

75. Zimmermann, M., et al., CRISPR screens identify genomic ribonucleotides as a source of PARP-trapping lesions. Nature, 2018. 559(7713): p. 285–289.

76. Olivieri, M. and D. Durocher, Genome-scale chemogenomic CRISPR screens in human cells using the TKOv3 library. STAR Protoc, 2021. 2(1): p. 100321.

77. Shannon, P., et al., Cytoscape: a software environment for integrated models of biomolecular interaction networks. Genome Res, 2003. 13(11): p. 2498–504.

78. Bindea, G., et al., ClueGO: a Cytoscape plug-in to decipher functionally grouped gene ontology and pathway annotation networks. Bioinformatics, 2009. 25(8): p. 1091–3.

79. Sanjana, N.E., O. Shalem, and F. Zhang, Improved vectors and genome-wide libraries for CRISPR screening. Nat Methods, 2014. 11(8): p. 783–784.

80. Brinkman, E.K., et al., Easy quantitative assessment of genome editing by sequence trace decomposition. Nucleic Acids Res, 2014. 42(22): p. e168.

81. Mojarad, B.A., et al., CEP19 cooperates with FOP and CEP350 to drive early steps in the ciliogenesis programme. Open Biol, 2017. 7(6).

82. McQuin, C., et al., CellProfiler 3.0: Next-generation image processing for biology. PLoS Biol, 2018. 16(7): p. e2005970.

83. Arganda-Carreras, I., et al., Trainable Weka Segmentation: a machine learning tool for microscopy pixel classification. Bioinformatics, 2017. 33(15): p. 2424–2426.

